# Disrupting miR-466l-3p and HuR Cooperation with Target Site Blockers Reveals a Therapeutic Strategy to Destabilize mRNA Transcripts

**DOI:** 10.64898/2026.03.30.709388

**Authors:** Vinod S. Ramgolam, Timur O. Yarovinsky, Sarah Huntenburg, Cheryl Bergman, Nancy Ruddle, Jeffrey R. Bender

## Abstract

MicroRNAs (miRNAs) typically regulate gene expression by promoting mRNA degradation, but select miRNAs, such as miR-466l-3p (miR-466), can instead stabilize transcripts in coordination with RNA-binding proteins (RBPs) like HuR. We identify conserved AU-rich elements (cAREs) within the 3′UTRs of *IL-17A*, *GM-CSF*, and *IL-23A* as critical *cis*-regulatory binding sites where miR-466 facilitates HuR recruitment to promote mRNA stability. Using site-directed mutagenesis, RNA pulldown, and MS2-TRAP assays to capture miRNA-mRNA complexes, we demonstrate that HuR binding depends on prior engagement by miR-466. Disrupting this interaction with rationally designed Target Site Blockers (TSBs) oligonucleotides destabilizes target mRNAs and suppresses cytokine expression *in vitro* and in *vivo*. TSBs directed against *IL-17A*, *GM-CSF*, and *IL-23A* selectively blocked miR-466 binding, reduced transcript stability, and lowered cytokine production without affecting unrelated mRNAs. In murine models of LPS-induced inflammation, psoriasis, and autoimmunity, TSBs exhibited therapeutic efficacy and cytokine specificity, outperforming monoclonal antibodies in some settings. Phosphorothioate-modified TSBs enabled systemic delivery and retained activity in human T cells, underscoring translational potential. Similar to antisense oligonucleotides, TSBs trigger RNase H1–mediated degradation while also blocking miRNA-mRNA interactions. These findings establish miR-466-HuR cooperation as a therapeutically targetable axis through TSBs without affecting global miRNA function.

**GRAPHICAL ABSTRACT:** 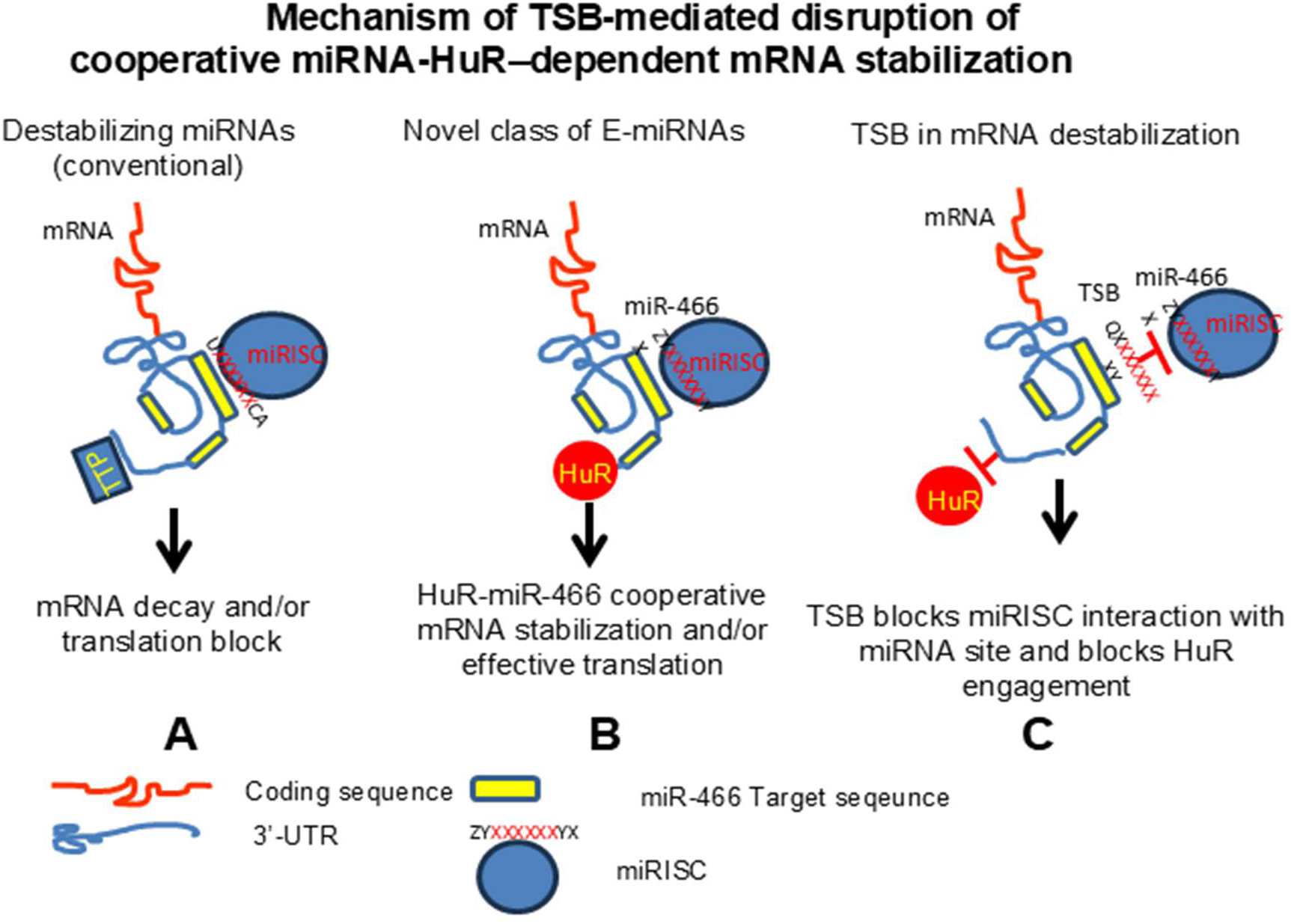

Mechanism of TSB-mediated disruption of cooperative miRNA-HuR–dependent mRNA stabilization
**A:** In the canonical model, destabilizing miRNAs (e.g., miR-16) bind to their target sites within the 3′UTR, recruiting the RNA-induced silencing complex (miRISC) to promote mRNA decay or translational repression.
**B:** In contrast, a newly identified class of miRNAs—stabilizing miRNAs (E-miRNAs), such as miR-466l-3p—bind to specific target sequences within AU-rich elements (AREs) in the 3′UTR. This binding facilitates cooperative recruitment of the RNA-binding protein HuR (ELAVL1), resulting in enhanced mRNA stability and/or translation.
**C:** Target site blockers (TSBs) designed to occlude miRNA-binding sites competitively inhibit miRISC loading, thereby disrupting HuR engagement and reversing stabilization. This selective disruption leads to transcript-specific mRNA destabilization without affecting global miRNA function.

## INTRODUCTION

The discovery of MicroRNAs (miRNAs) by Drs. Victor Ambros and Gary Ruvkun in the early 1990s, for which they were awarded the 2024 Nobel Prize in Medicine, revolutionized our understanding of gene regulation^1,2^. miRNAs are small non-coding RNAs that bind complementary sequences within target mRNAs-typically in the 3′ untranslated region (3′UTR)-to modulate mRNA stability and translation. Despite the therapeutic potential of miRNA mimics and inhibitors (anti-MIRs), clinical translation has been limited by issues of target specificity, off-target effects, and delivery challenges. Despite over 30 clinical trials, no miRNA-targeted therapy has yet received FDA approval^3^.

miRNAs function within a complex post-transcriptional regulatory landscape that includes *cis*-acting elements, such as AU-rich elements (AREs), and trans-acting factors like RNA-binding proteins (RBPs)^4,5^. AREs are conserved motifs that mediate rapid mRNA turnover and are enriched in transcripts encoding cytokines and immune regulators^6^. RBPs such as HuR (ELAVL1) and TTP (ZFP36) bind to AREs to promote either stabilization or decay^7–9^. While miRNAs have long been considered independent effectors of post-transcriptional regulation, emerging evidence suggests they often function in coordination or in competition with RBPs to fine-tune gene expression or when dysregulated in disease pathology^10^. There are several studies indicating that HuR and miRNAs exert antagonistic^10,11^ or cooperate in their post-transcriptional regulatory function^12,13^. For instance, HuR can antagonize the actions of miR-200, miR-181, and miR-637 by binding near their target sites on mRNAs^14–16^. Conversely, some RBPs may cooperate with miRNAs to stabilize target transcripts, as demonstrated by the synergistic action of HuR and miR-3134^17^.

We previously showed that β2-integrin LFA-1 engagement with intercellular adhesion molecule-1 (ICAM-1) on lymphocytes induces HuR-mediated stabilization of cytokine mRNAs, promoting inflammatory responses^8,9,18,19^. However, miRNAs such as mmu-miR-466l-3p (hereafter miR-466), which target ARE-like sequences, may cooperate with or antagonize RBPs like HuR. miR-466 is expressed in both mouse and human cells and has been reported to stabilize IL-10 mRNA or upregulate other pro-inflammatory cytokines while promoting degradation of IFN-α transcripts, depending on cellular context^20–22^. These opposing functions suggest that miR-466 may influence the dynamic regulation of RBP–mRNA interactions.

We hypothesized that miR-466 and HuR co-regulate ARE-containing mRNAs involved in Th17-mediated inflammation through a cooperative mechanism. Among the key transcripts of interest—IL-17A, GM-CSF, and IL-23A—each contains conserved AREs and is critically regulated post-transcriptionally. These cytokines form the core of the Th17 axis implicated in autoimmune and inflammatory diseases including multiple sclerosis, IBD, psoriasis, and rheumatoid arthritis^23–29^. Collectively, these transcripts form a tightly regulated inflammatory axis with potent pathogenic potential when dysregulated, and together represent validated and evolving targets for RNA- and antibody-based therapies in immune-mediated disease^30–33^.

Here, we define a previously unrecognized cooperative interaction between miR-466 and HuR at AREs within the 3′UTRs of IL-17A, GM-CSF, and IL-23A. Employing site directed mutagenesis, RNA-IP, and miRNA pull-down assays, we demonstrate that miR-466 facilitates HuR binding and transcript stabilization. We further show that this interaction can be selectively disrupted by Target Site Blocker (TSB) oligonucleotides, which block miR-466 access to its binding sites, leading to mRNA destabilization and cytokine suppression. These findings advance our understanding of miRNA–RBP synergy and introduce a tractable therapeutic strategy for selective cytokine inhibition in Th17-driven diseases.

## RESULTS

### Role of HuR in IL-17A mRNA Stability and Expression in T Cells and EAE Pathogenesis

To investigate the role of HuR in human IL-17A mRNA stability, we used AREsite (http://nibiru.tbi.univie.ac.at/cgi-bin/AREsite/AREsite.cgi) to identify multiple AU-rich elements (AREs) within the human IL-17A 3′UTR. We first assessed HuR binding in primary human T cells stimulated with low-dose (2 ng/ml) phorbol myristate acetate (PMA) to activate integrins and gene transcription followed by plating on recombinant human ICAM-1(rhICAM)– or poly-L-lysine (PLL)–coated surfaces. HuR-RNA immunoprecipitation (RIP) revealed a ∼3,000-fold enrichment of IL-17A mRNA in ICAM-1–adherent cells, indicating integrin-dependent HuR binding (Fig. 1A). To assess IL-17A mRNA stability, we induced transcriptional arrest with DRB (dichlorobenzimidazole 1-β-D-ribofuranoside) treatment to inhibit RNA synthesis. Total RNA was harvested at defined intervals and IL-17A transcript levels were quantified by qRT-PCR, normalized to the initial time point (T = 0 min). In T cells adhered to ICAM-1, IL-17A mRNA levels remained essentially unchanged over the 60-minute assay, indicating complete stabilization. In contrast, cells plated on PLL exhibited rapid mRNA decay, with IL-17A transcript levels declining to ∼10% of baseline within 30 minutes (Fig. 1B).

**Figure 1:**
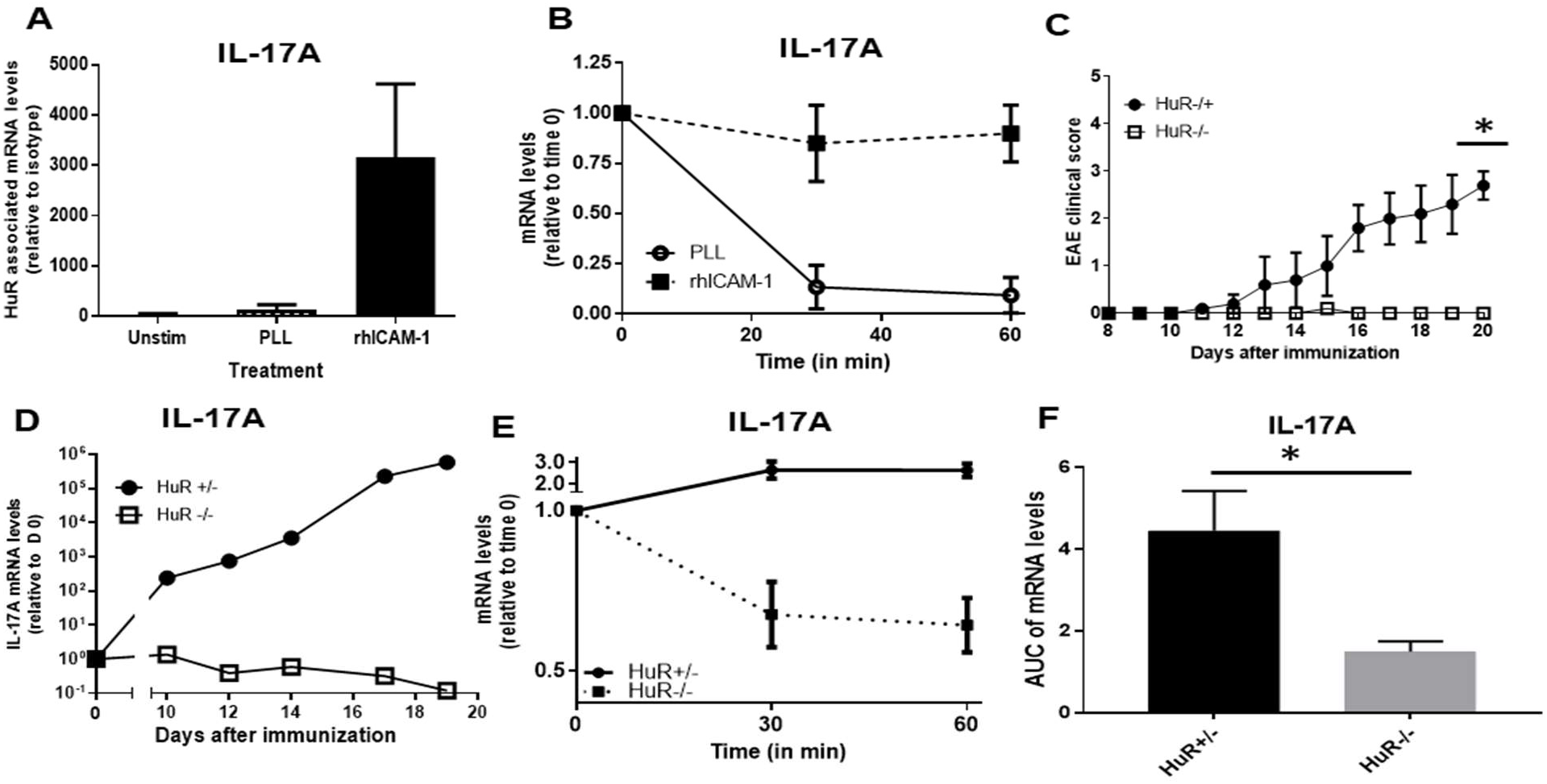
HuR regulates IL-17A mRNA stability through integrin engagement and post-transcriptional control in human T cells and EAE models. ***(A*)** Primary human peripheral T cells were either left unstimulated or stimulated with 2 ng/mL PMA for 2 hours on plates coated with either poly-L-lysine (PLL) or rhICAM-1. Following stimulation, cells were crosslinked, lysed, and immunoprecipitated with anti-HuR or IgG1 control antibodies. Bound IL-17A and GAPDH mRNAs were quantified by qRT-PCR, and enrichment was calculated relative to IgG1 controls. (*B*) Primary human T cells were stimulated as described above, and transcription was inhibited after two hours with 0.2 mM DRB. Total RNA was harvested at the indicated time points, and IL-17A and GAPDH mRNA levels were quantified by qRT-PCR. Fold changes were calculated relative to transcript levels at time zero (T = 0), corresponding to the point of transcriptional inhibition. (*A,B*), Data represent three independent experiments, with all samples analyzed in triplicate and presented as mean ± standard error of the mean (S.E.M. (**C)** EAE was induced in HuR⁻/⁻ (n=5) and HuR+/⁻ (n=5) mice by immunization with 200 µg MOG₃₅–₅₅ peptide emulsified in 250 µg CFA. Clinical symptoms were assessed daily. Two-way ANOVA revealed statistically significant differences between groups on days 19 and 20. (*D*) One mouse per group was euthanized at days 0, 10, 12, 14, 17, and 19 for CNS collection. Lumbar spinal cords were isolated and analyzed by qRT-PCR for IL-17A mRNA. Fold changes were calculated relative to naïve (day 0) controls. (*E*) Splenic T cells from HuR⁺/⁻ and HuR⁻/⁻ mice were harvested at day 21 post-EAE induction and cultured with 10 µg/mL MOG₃₅–₅₅ for 5 days. Cells were restimulated with PMA and plated on ICAM-1– or PLL-coated wells. Total RNA was collected at indicated time points for IL-17A mRNA decay analysis. (*F*) A Welch’s t-test was performed to compare the AUC of IL-17A mRNA levels between HuR+/- and HuR-/- groups. Data are from three independent experiments, each performed in triplicate. Results are shown as mean ± SEM. (*) *P* < 0.05 was considered statistically significant.

To assess the *in vivo* relevance, we crossed HuR^fl/fl^ ^19^mice with RORγt-Cre ^34^mice to generate T cell–specific HuR knockouts (HuR⁻/⁻). We induced experimental autoimmune encephalomyelitis (EAE) using the MOG₃₅–₅₅ peptide and complete Freunds Adjuvant (CFA) in both HuR^⁻/⁻^ and control HuR^+/-^ mice. Clinical symptoms were monitored and scored over a 20-day period. Strikingly, HuR^⁻/⁻^ mice failed to develop any clinical signs of EAE, whereas HuR^+/⁻^ mice exhibited typical disease progression (Fig. 1C). Chen et al. used OX40-Cre mice to delete HuR expression, but they observed less pronounced effects compared to those seen with RORγt-Cre–mediated deletion^35^. In a parallel experimental setup spinal cords collected from immunized mice at days 0, 10, 12, 14, 17 and 19 revealed a marked increase in IL-17A mRNA in HuR⁺/⁻ mice, rising from ∼240-fold at day 10 to ∼500,000-fold by day 19, while IL-17A remained nearly undetectable in HuR⁻/⁻ mice (Fig. 1D). GM-CSF mRNA followed a similar trend and is shown in Fig. S1A; TNF-α, IFN-γ, and RORγt also exhibited comparable kinetics (data not shown). To assess HuR’s role in IL-17A mRNA stability, we performed decay assays on MOG₃₅-₅₅ expanded splenic lymphocytes from HuR⁻/⁻ and HuR⁺/⁻ mice. Following ICAM-1 adhesion and 2 ng/mL PMA stimulation, transcription was halted with DRB. IL-17A mRNA remained stable in HuR⁺/⁻ cells but decayed ∼35% in HuR⁻/⁻ T cells and was statistically significant through area under the curve analysis (AUC) (Fig. 1E&F). These findings demonstrate that LFA-1 integrin engagement enhances HuR-dependent stabilization of IL-17A mRNA in T cells, and that HuR deficiency abrogates Th17-mediated neuroinflammation—establishing HuR as a key post-transcriptional regulator of IL-17A and a critical mediator of the Th17 axis during EAE.

### Conserved AU-Rich Element in the IL-17A 3′UTR Control mRNA Stability

The human IL-17A 3′UTR (1,347 nt) contains three conserved AU-rich elements (cAREs) clustered within a 621-nt region, with a 32-bp segment showing high interspecies conservation (Fig. 2A). To investigate the role of AREs in IL-17A mRNA post-transcriptional regulation, we used the pBBB β-globin reporter system, which expresses a highly stable rabbit β-globin transcript under a serum-inducible *c-fos* promoter and enables assessment of regulatory UTR elements on mRNA decay. We cloned either the IL-17A full-length 3′UTR (WT), the ARE-containing fragment (ARE), or the distal non-ARE region (NARE) into pBBB and transfected these into NIH-3T3 cells along with an eGFP plasmid for normalization. While β-globin mRNA remained stable with pBBB and pBBB-17-NARE constructs, transcripts harboring either the 17-WT 3′UTR or the 17-ARE fragment exhibited rapid decay, confirming that the AREs are sufficient to destabilize mRNA (Fig. 2B). To assess translational impact, we cloned the IL-17-WT downstream of the firefly luciferase gene in the pMIRGlo system (pMIR-17-WT), and observed reduced luciferase expression in transfected cells compared to empty vector control. This suppression was reversed by anisomycin, a p38 MAPK activator known to promote HuR cytoplasmic translocation, suggesting that HuR counteracts ARE-mediated decay (Fig. S2).

**Figure 2:**
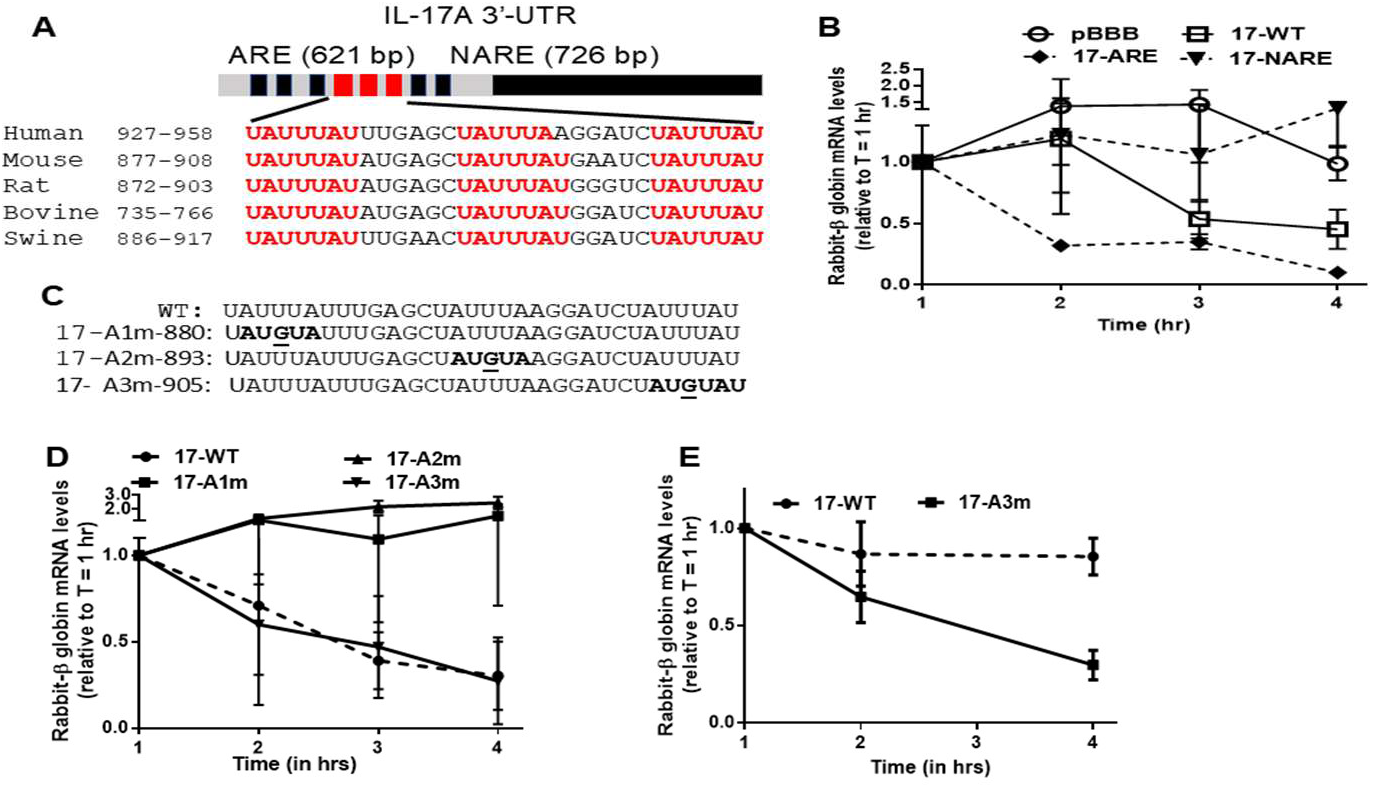
Characterization of IL-17A mRNA 3’-UTR and Its Role in Post-Transcriptional Regulation. **(*A***) The IL-17A 3′UTR contains a conserved ARE region over several species (621 bp) and three clustered cAREs within a 32-bp conserved stretch. **(*B***) The full-length IL-17A 3′UTR (WT), as well as the ARE and NARE subfragments, were cloned into the pBBB reporter plasmid containing a rabbit β-globin gene and co-transfected with an enhanced GFP (eGFP) plasmid into NIH-3T3 cells for normalization. Following overnight serum starvation, cells were stimulated with 20% serum, and RNA was harvested at 1, 2, 3, and 4 hours post-stimulation. β-globin and GFP mRNA levels were quantified by qRT-PCR, with β-globin levels normalized to GFP and expressed as fold change relative to the 1-hour time point. **(*C***) Point mutations were introduced into the first (17-A1m), second (17-A2m), and third (17-A3m) conserved AREs within the IL-17A 3′UTR in the pBBB-17-WT construct. **(*D*)** NIH-3T3 cells were transfected with WT or cARE-mutant constructs and subjected to mRNA decay analysis as in (B). **(*E*)** Jurkat T cells were transfected with WT or 17-A3m reporters and stimulated via ICAM-1 adhesion; β-globin mRNA stability was assessed post-serum stimulation

To further dissect the roles of the individual cAREs, we introduced point mutations (AT**T**TA → AT**G**TA) at each the 3 cAREs within the pBBB-17-WT construct, generating pBBB-17-A1m, -17-A2m, and -17-A3m (Fig. 2C). Point mutations were chosen to avoid the potential for multiple sequence changes to drastically alter RNA folding and disrupt the function of the targeted regio. mRNA decay assays in NIH-3T3 cells revealed that mutations in the first or second cAREs did not significantly affect β-globin mRNA stability, whereas mutation of the third cARE recapitulated the decay kinetics of the WT construct, identifying it as a dominant destabilizing element (Fig. 2D). To validate the role of cARE3 in LFA-1 induced HuR-dependent mRNA stabilization, Jurkat T cells were transfected with either pBBB-17-WT or the mutant pBBB-17-A3m and subjected to ICAM-1–mediated adhesion and serum stimulation. While the WT transcript remained stable, the cARE3 mutant exhibited enhanced decay with ∼ 25% remaining after 4 hrs, indicating that cARE3 is essential for LFA-1/ICAM-1–induced mRNA stabilization (Fig. 2E). These results identify cARE3 as the dominant regulatory element within the IL-17A 3′UTR and a critical role for HuR-dependent transcript stabilization.

### miR-466 as a Key Post-Transcriptional Regulator of IL-17A Expression through ARE-Dependent Stabilization and HuR Interaction

miR-466 is a microRNA that recognizes ARE-like motifs. Its seed sequence (AUAAAUA) overlaps with AUUUA-containing elements found within cAREs in the IL-17A 3′UTR. To clarify prior inconsistencies in nomenclature, we refer to the mature miRNA sequence (5′-UAUAAAUACAUGCACACAUAUU-3′) as miR-466 throughout, corresponding to mmu-miR-466l-3p and its nearly identical human ortholog (Fig. 3A). The human miR-466 is encoded within a genomic region on chromosome 4^21^. To assess the impact of miR-466 on IL-17A mRNA decay, we transfected the pBBB-17A-WT construct into NIH-3T3 cells stably overexpressing miR-466 (466-3T3) or a scrambled control (SC-3T3), and compared transcript stability between the two cell lines. The rabbit β-globin transcripts remained stable in 466-3T3 cells but decayed rapidly in SC-3T3, indicating that miR-466 is involved in iL-17A mRNA stabilization (Fig. 3B).To confirm direct binding of miR-466 to the IL-17A 3’UTR, we used the MS2-tagged RNA affinity purification (MS2-TRAP) pull-down assays ^36^. This system employs a GST pull-down assay to isolate miRNA-mRNA complexes and assess direct binding. WT-3T3, SC-3T3, and 466l-3T3 cells were co-transfected with pMS2-17-WT and pMS2-GST plasmids. As a control for non-specific miRNA binding, cells were also transfected with empty pMS2 and pMS2-GST plasmids. miR-466 enrichment in the pull-down fraction was ∼6–7-fold in WT and SC cells, and ∼53-fold in miR-466–overexpressing cells, confirming specific binding to the IL-17A 3′UTR (Fig. 3C).

**Figure 3.**
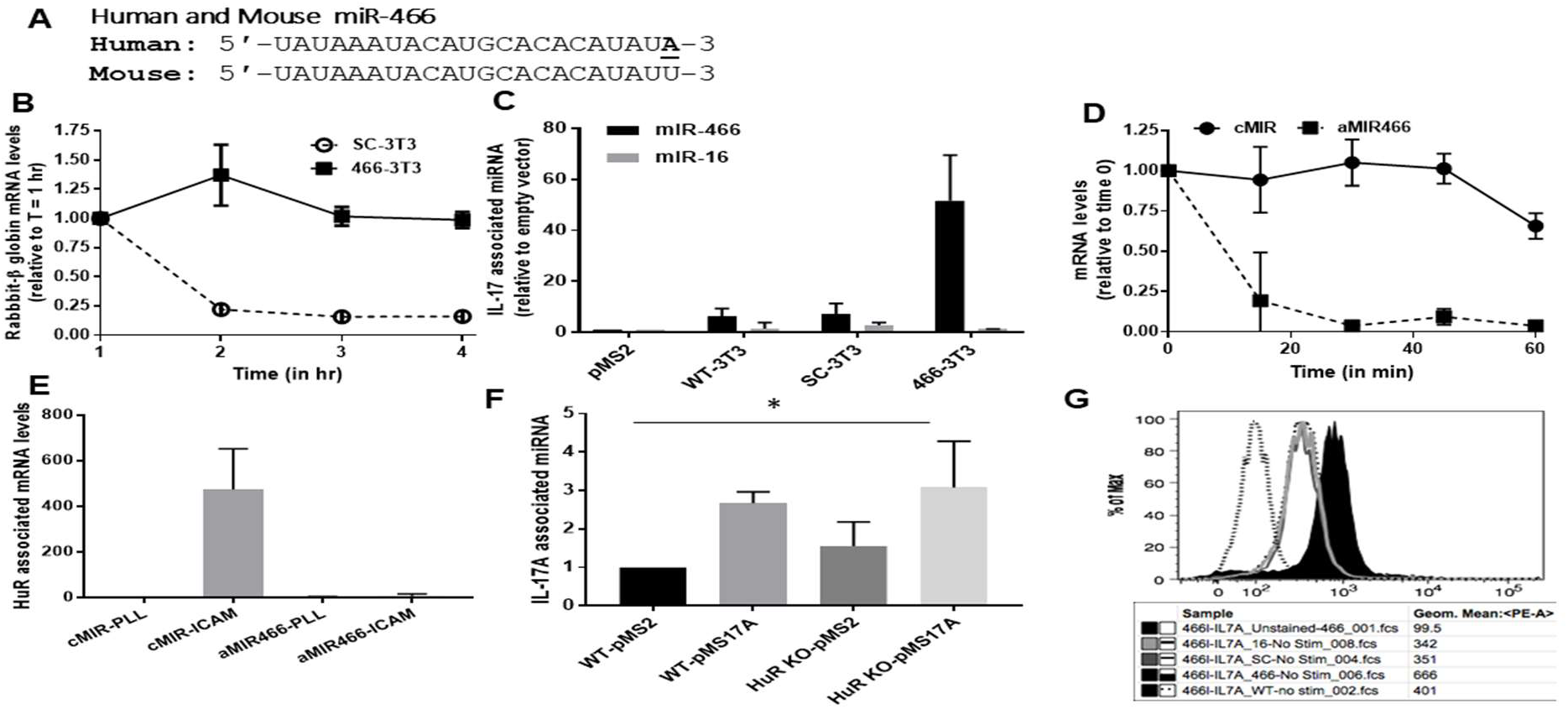
miR-466 as a Key Post-Transcriptional Regulator of IL-17A Expression through ARE-Dependent Stabilization and HuR Interaction. (*A*) Sequence alignment of human and mouse miR-466. (*B*) mRNA decay assay using rabbit β-globin reporter in SC-3T3 and miR-466–overexpressing (466-3T3) cells transfected with pBBB-h17A. Following serum induction for 1 hour, β-globin transcript levels were measured at the indicated time points by qRT-PCR as described earlier. (*C*) MS2-TRAP assay performed in NIH-3T3, SC-3T3, and 466-3T3 cells co-transfected with either the empty pMS2 vector or pMS2 containing the IL-17A 3′UTR (pMS2-17-WT), along with pMS2-GST. Associated miR-466 and miR-16 levels were quantified by qRT-PCR, normalized to U6 snRNA, and expressed as fold enrichment relative to the empty pMS2 control. (*D*) Primary human T cells were transfected with 25 nM of either an anti-miR-466 inhibitor (aMIR466) or a scrambled control oligonucleotide (cMIR) were subjected to LFA-1 mediated mRNA stabilization as described before. (*E*), Primary human peripheral T cells were transfected and stimulated as in *D*. After 2 hours, cells were cross-linked, lysed, and subjected to HuR-IP. Associated IL-17A mRNA levels were quantified by qRT-PCR and normalized to input. Data represent mean ± SEM from three independent experiments. (*F*) MS2-TRAP was performed in HuR⁺/⁻ and HuR⁻/⁻ mouse T cells transfected with pMS2-17-WT or empty vector plus pMS2-GST. After 2 hours on ICAM-1–coated plates, lysates underwent GST pull-down, and miR-466 and U6 levels were quantified by qRT-PCR relative to vector control. Data are presented as mean ± S.E.M. from three independent experiments. Statistical significance was determined using one-way ANOVA. (*) *P* < 0.05 was considered significant. (*G*) Flow cytometry of IL-17A in EL-4 cells stably expressing miR-466, miR-16, or SC, with or without PMA/ionomycin stimulation for 6 hours. Intracellular IL-17A was measured as geometric mean fluorescence intensity (gMFI)

To evaluate miR-466’s role LFA-1/HuR dependent IL-17A mRNA stabilization, human T cells were transfected with a miR-466 inhibitor (aMIR466) or control anti-miR (cMIR), then subjected to LFA-1 engagement. Inhibition of miR-466 led to IL-17A mRNA decay while cMIR preserved transcript stability (Fig. 3D). We then determined if miR-466 affects the association of HuR with IL-17A mRNA, we performed HuR-RNA immunoprecipitation assays. Human primary T cells obtained as described above and transfected with either aMIR466 or cMIR were plated on PLL or ICAM-1, crosslinked, lysed by sonication, and subjected to immunoprecipitation with anti-HuR or isotype control antibodies. HuR-RIP assays revealed a ∼300-fold increase in HuR binding to IL-17A mRNA in ICAM-1–adhered cells transfected with cMIR, whereas HuR association remained at baseline in cells treated with aMIR466 (Fig. 3E). Thus, miR-466 is required for HuR association with IL-17A mRNA for it’s stabilization.

The reciprocal effect of HuR on miR-466 binding to IL-17A mRNA was further investigated in HuR deleted T cells. We conducted pull-down assays in HuR⁺/⁻ and HuR⁻/⁻ mouse T cells transfected with the pMS2-17-WT construct. Following transfection, cells were adhered to ICAM-1 for 2 hours to mimic LFA-1 induced HuR mRNA association and then subjected to GST pull-down assays. Compared to the empty vector, miR-466 showed increased binding to the pMS2-17-WT construct in comparison to the control pMS2. However, this binding was equivalent in both HuR⁺/⁻ (∼2.6-fold) and HuR⁻/⁻ (∼2.7-fold) cells (Fig. 3F), indicating that miR-466 associates with IL-17A mRNA independently of HuR. The absence of differences in binding between HuR⁺/⁻ and HuR⁻/⁻ cells implies that HuR is not essential for miR-466 recruitment to IL-17A mRNA. This was in line with earlier HuR-RIP experiments demonstrating that HuR requires miR-466 for robust association with IL-17A mRNA. We next tested the impact of miR-466 on IL-17A protein expression. EL-4 cells stably expressing miR-466 and stimulated for cytokine expression with PMA/Ionomycin exhibited significantly increased IL-17A expression by intracellular flow cytometry (gMFI = 666 vs. 401 in control cells) (Fig. 3G), indicating that miR-466 enhances IL-17A expression. Together, these findings demonstrate that miR-466 directly binds to the IL-17A 3′UTR, stabilizes its transcript via cAREs, and promotes HuR recruitment and protein expression.

### The blockade of miR-466 binding to IL-17A 3’UTR with site specific TSB prevents IL-17A mRNA stabilization and expression

To map the specific miR-466 target site within the IL-17A 3′UTR, we performed MS2-TRAP pull-down assays using constructs harboring point mutations in each of the three cARE1–3 (Fig. 2A). The resulting constructs—pMS2-17-A1m, pMS2-17-A2m, and pMS2-17-A3m—along with the WT and empty pMS2 vectors, were co-transfected into NIH-3T3 cells with the pMS-GST plasmid, and miRNA pull-down assays were performed. Consistently, miR-466 binding was enriched ∼2–3-fold in cells transfected with pMS2-17-WT, -A1m, or -A2m constructs, but was markedly diminished to baseline levels in the 17-A3m mutant (Fig. 4A). Mutation of cARE3 (17-A3m) significantly reduced miR-466 binding, while mutations in cARE1 (17-A1m) or cARE2 (17-A2m) had no effect, identifying cARE3 as the dominant miR-466 interaction site. Initial MS2-TRAP assays were performed in wild-type cells, as stable miRNA overexpression could disrupt normal cellular function and confound experimental results.

**Figure 4.**
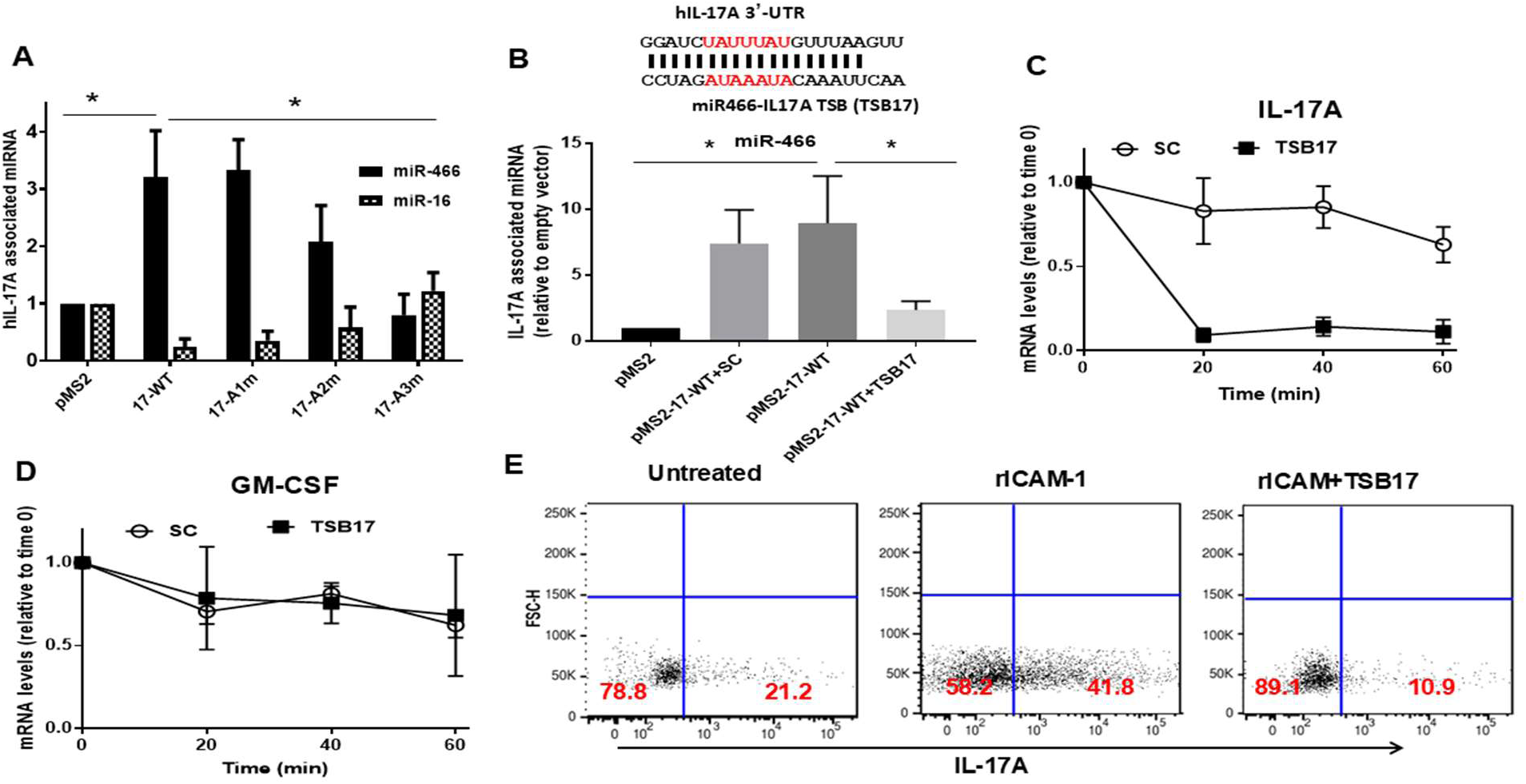
TSB17 selectively blocks miR-466 interaction with the IL-17A 3′UTR and disrupts cytokine expression in vitro and in vivo. **(A)** NIH-3T3 cells were co-transfected with pMS2-GST and either empty pMS2, pMS2-IL-17-WT, or cARE-mutant constructs for the MS-TRAP assay. miR-466 and miR-16 enrichment was quantified by qRT-PCR and normalized against U6. **(*B*)** Sequence alignment of the miR-466 binding site in the IL-17A 3′UTR with complementary TSB17. 466-3T3 cells were transfected with pMS2-IL-17-WT ± SC or TSB17, followed by crosslinking and MS2-TRAP. Bound miR-466 was quantified by qRT-PCR. (***A, B)*** Statistical significance was determined using two-way ANOVA. (*) *P* < 0.05 was considered significant. (***C, D***) Human primary T cells (∼1×10⁶) were transfected with TSB17 or SC, stimulated with PMA, and plated on ICAM-1–coated surfaces. After transcriptional arrest with DRB, IL-17A (***C***) and GM-CSF (***D***) mRNA levels were measured by qRT-PCR over time to assess stabilization. **(*E*)** T cells from MOG₃₅–₅₅–immunized 2D2-transgenice mice were expanded with peptide, electroporated with 25 nM TSB17, and stimulated with PMA on ICAM-1–coated plates. After 6 hours, cells were fixed, stained for IL-17A, and analyzed by flow cytometry. Data represent one of three independent experiments.

To specifically block miR-466 interaction at target site, we designed a Target Site Blocker (TSB17) complementary to the third cARE, spanning the core target site and including 3–4 nucleotides on the 5’ and 3’ flanking sites (Fig. 4B). MS2-TRAP assays in NIH-3T3 cells overexpressing miR-466 (466-3T3) showed that transfection with the pMS2-17-WT construct led to an ∼8-fold increase in miR-466 binding compared to the empty vector. Co-transfection with TSB17, but not scramble control (SC), significantly reduced miR-466 binding, confirming that TSB17 specifically blocks miR-466 interaction with the IL-17A 3′UTR (Fig. 4B). To evaluate the functional impact of TSB17, we transfected anti-CD3 (OKT3)-expanded human T cells with TSB17 or SC oligos, then performed mRNA stabilization assays following LFA-1 engagement. TSB17 caused rapid IL-17A transcript decay to ∼ 10 % at time 20 mins, while SC-treated cells ∼ 75 % retained mRNA stability over 60 mins (Fig. 4C). GM-CSF mRNA, also a miR-466 target, remained unaffected, indicating that TSB17 selectively blocks miR-466–IL-17A interactions without depleting miR-466 availability (Fig. 4D). This specificity is critical to minimize off-target effects and preserve endogenous miRNA regulatory networks, reinforcing the therapeutic value of TSB strategy.

To further assess the impact of TSB17 on IL-17A protein expression, we employed the dual-luciferase pMIRGlo reporter system. NIH-3T3 cells were transfected with pMIR-17A-WT in the presence or absence of TSB17 or SC oligonucleotides. Cells transfected with TSB17 exhibited significantly lower firefly luciferase activity compared to SC or vector controls, consistent with reduced transcript stability and diminished protein translation (Fig. S3). To validate in TSB17 in primary cells, we used T cells derived from MOG₃₅–₅₅–immunized mice. Splenocytes from EAE mice at day 21 after immunization were stimulated 10 ug/ml MOG₃₅–₅₅ peptide for 5 days. The cells were transfected with TSB17 followed by adhesion to ICAM-1. ICAM-1 adhesion induced IL-17A in 41.8% of untreated cells (vs. 21.2% in baseline). TSB17 reduced IL-17A+ cells to 10.9%, confirming the blockade of integrin-mediated transcript stabilization (Fig. 4F). These results demonstrate that TSB17 selectively prevents miR-466 binding to IL-17A mRNA, resulting in transcript destabilization and suppression of IL-17A protein expression.

### *In Vivo* Evaluation of TSB17 Across Models of Inflammation

To evaluate the therapeutic potential of TSB17, we tested it in four IL-17A–driven inflammation models: LPS-induced systemic inflammation, imiquimod-induced psoriasis, EAE, and experimental autoimmune uveitis (EAU). For the LPS cytokine storm model, we used C57BL/6 mice and IL-17A knockout mice^37^ which received intraperitoneal TSB17 or SC oligo 16 hours before and at the time of LPS challenge. Serum IL-17A levels measured 7 hours post-LPS were nearly undetectable in TSB17-treated mice, comparable to IL-17A-knockouts, while SC-, and LPS treated only mice exhibited ∼200 pg/mL IL-17A (Fig. S4A). IL-6 levels, also miR-466–regulated, remained unaffected (Fig. S4B), confirming the specificity of TSB17 for IL-17A. In the imiquimod-induced psoriasis model, C57BL/6 and IL-17A knockout mice received daily topical imiquimod and were treated with SC or TSB17 formulated in 10% Pluronic F-127 hydrogel. Disease severity, scored using the Psoriasis Area and Severity Index (PASI), revealed that TSB17-treated mice exhibited significantly reduced erythema and scaling compared to control-treated mice (Fig. S4C). TSB17 significantly reduced erythema and scaling (Fig. 5A), indicating effective topical suppression of IL-17A–mediated inflammation. To evaluate the efficacy of TSB17 in an autoimmune setting, we used a zymosan-adjuvanted EAE model known to be highly dependent on IL-17A signaling. 2D2-TCR transgenic mice immunized with MOG₃₅–₅₅ and zymosan developed severe EAE symptoms, whereas 2D2-TCR-IL-17A knockout mice remained largely disease-free, confirming the model’s IL-17A dependency (Fig. S4D). In prophylactic studies, TSB17 was administered at 5 mg/kg starting on day 6 post-immunization, followed by 10 mg/kg on day 9 and single 5 mg/kg doses every three days. This regimen completely prevented disease onset up to 36 days, while SC-treated mice developed severe symptoms requiring euthanasia by day 20 (Fig. 5B). In therapeutic studies, TSB17 was administered after mice reached a clinical score of 1 and effectively halted disease progression, whereas SC-treated animals continued to worsen. Notably, TSB17 outperformed anti–IL-17A neutralizing antibodies treatment, highlighting a potential mechanistic advantage of TSB penetration to the CNS (Fig. 5C).

**Figure 5:**
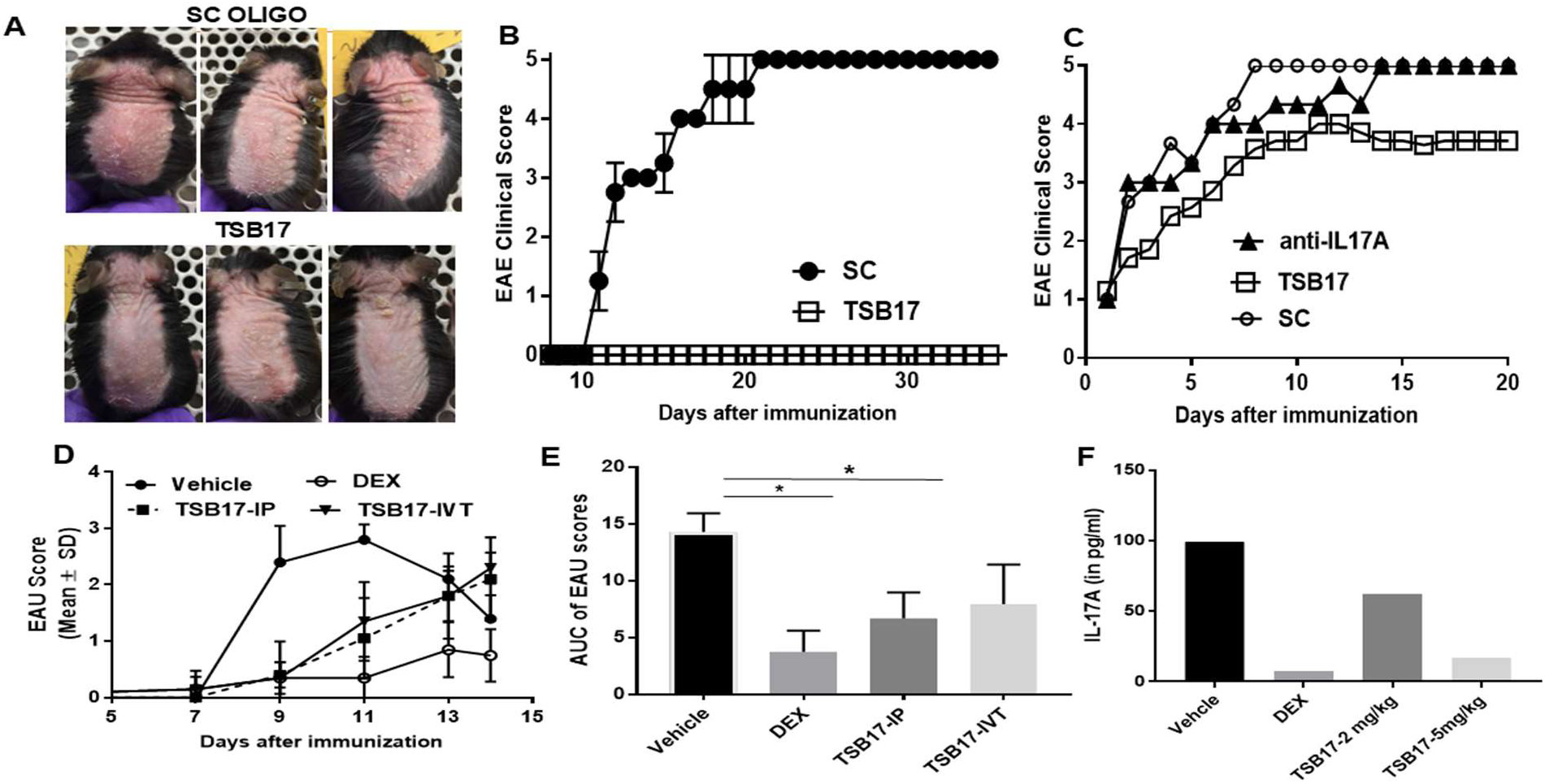
Therapeutic efficacy of TSB17 in IL-17A–mediated inflammatory animal models. (***A*)** C57BL/6 mice were treated daily with imiquimod for 5 days to induce psoriasis-like inflammation and received topical TSB17 or SC oligonucleotides (n=3/group) in 10% Pluronic F-127, applied 8 hours post-imiquimod. (***B***) 2D2-TCR transgenic mice were immunized subcutaneously at two sites with 100 μL of an emulsion containing 250 μg MOG₃₅–₅₅ peptide and 500 μg zymosan per mouse. Mice were administered intraperitoneally (IP) TSB17 (n=5) or SC oligonucleotide (n=4) at 5 mg/kg on day 6, followed by 10 mg/kg on day 9, and 5 mg/kg every three days thereafter. Clinical EAE scores were recorded daily. (***C***) 2D2-TCR transgenic mice were immunized as above with MOG₃₅–₅₅ + zymosan–induced EAE were monitored until they reached a clinical score of 1, at which point treatment was initiated. Mice were administered either daily doses of 5 mg/kg TSB17 (n=7), SC oligonucleotide (n=3), or every other day of 100 μg/mouse of anti–IL-17A neutralizing antibody (n=3). **(*D*)** EAU was induced in healthy rats (n=5 per group), which were then either left untreated, treated with intravitreal (IVT) dexamethasone (40 µg on days 6 and 9), or administered IL-17A TSB either IP (5 mg/kg daily from days 6–10) or IVT (120 µg on days 6 and 9). EAU scores were recorded at specified time points. (***E***) Area under the curve (AUC) analysis was performed for each cohort, with statistical significance assessed by one-way ANOVA (*P < 0.005). (***F***) Aqueous humor from five rats per treatment group (TSB17 or DEX) was pooled and analyzed for IL-17A protein levels.

We also investigated the therapeutic potential of a rat-specific TSB17 in the EAU model. Rats were immunized with interphotoreceptor retinoid-binding protein (IRBP) in CFA and treated with TSB17 via both intravitreal and intraperitoneal routes. Clinical EAU scores were assessed on days 0, 7, 9, 11, 13, and 14 post-immunization. TSB17-treated rats exhibited a significant reduction in EAU severity compared to vehicle-treated controls, with average scores reduced from ∼3 (severe inflammation) to ∼1 (mild inflammation). Notably, intravitreal dexamethasone (40 µg), a standard corticosteroid, reduced EAU severity by approximately 75%, while systemic administration of TSB17 achieved around a 50% reduction (Fig. 5D). AUC analysis confirmed that IP delivery of TSB17 significantly lowered EAU scores (Fig. 5E). We also evaluated the impact of TSB17 treatment on IL-17A levels in the aqueous humor (AH). AH samples from all five rats per group were pooled for analysis. In SC-treated rats, IL-17A levels averaged 99.01 pg/mL. TSB17 treatment resulted in a dose-dependent reduction, with levels decreasing to 62.32 pg/mL at 2 mg/kg and 16.79 pg/mL at 5 mg/kg. For comparison, dexamethasone-treated rats exhibited IL-17A levels of 7.6 pg/mL (Fig. 5F). Together, these studies demonstrate that TSB17 is effective in suppressing IL-17A–driven inflammation in systemic, cutaneous, and autoimmune settings.

### Post-transcriptional regulation of GM-CSF mRNA by HuR and miR-466 in Th17-mediated inflammation

HuR stabilizes GM-CSF mRNA during inflammatory responses, a process further enhanced by integrin-mediated adhesion—specifically LFA-1 binding to ICAM-1—which activates intracellular signaling pathways that promote the stabilization of labile cytokine transcripts^8,19^. The GM-CSF 3′UTR contains multiple cAREs overlapping predicted miR-466 seed sequences, suggesting a similar mechanism of regulation. Given the shared regulatory features of IL-17A and GM-CSF, we examined whether miR-466 and HuR also cooperate to stabilize GM-CSF mRNA.

We introduced point mutations at seven predicted cAREs and one non-target control site within the GM-CSF 3′UTR (Fig. 6A). Wild-type (pMS2-GM-WT) and mutant constructs (pMS2-GM-A1m to -A7m, with CNT serving as a negative control) were co-transfected with pMS2-GST into NIH-3T3 cells, and miRNA pull-down assays were performed using the MS2-TRAP system. Most constructs exhibited 7–8-fold enrichment of miR-466 binding relative to vector control. In contrast, binding was reduced to ∼2-fold for the GM-A6m and GM-A8m mutants, while mutation of GM-A4m resulted in the most pronounced loss of binding to ∼ 1 fold, identifying cARE4 as the dominant miR-466 regulatory site in the GM-CSF 3′UTR (Fig. 6B). We also measured the binding of miR-16, which was not affected in the different samples.

**Figure 6:**
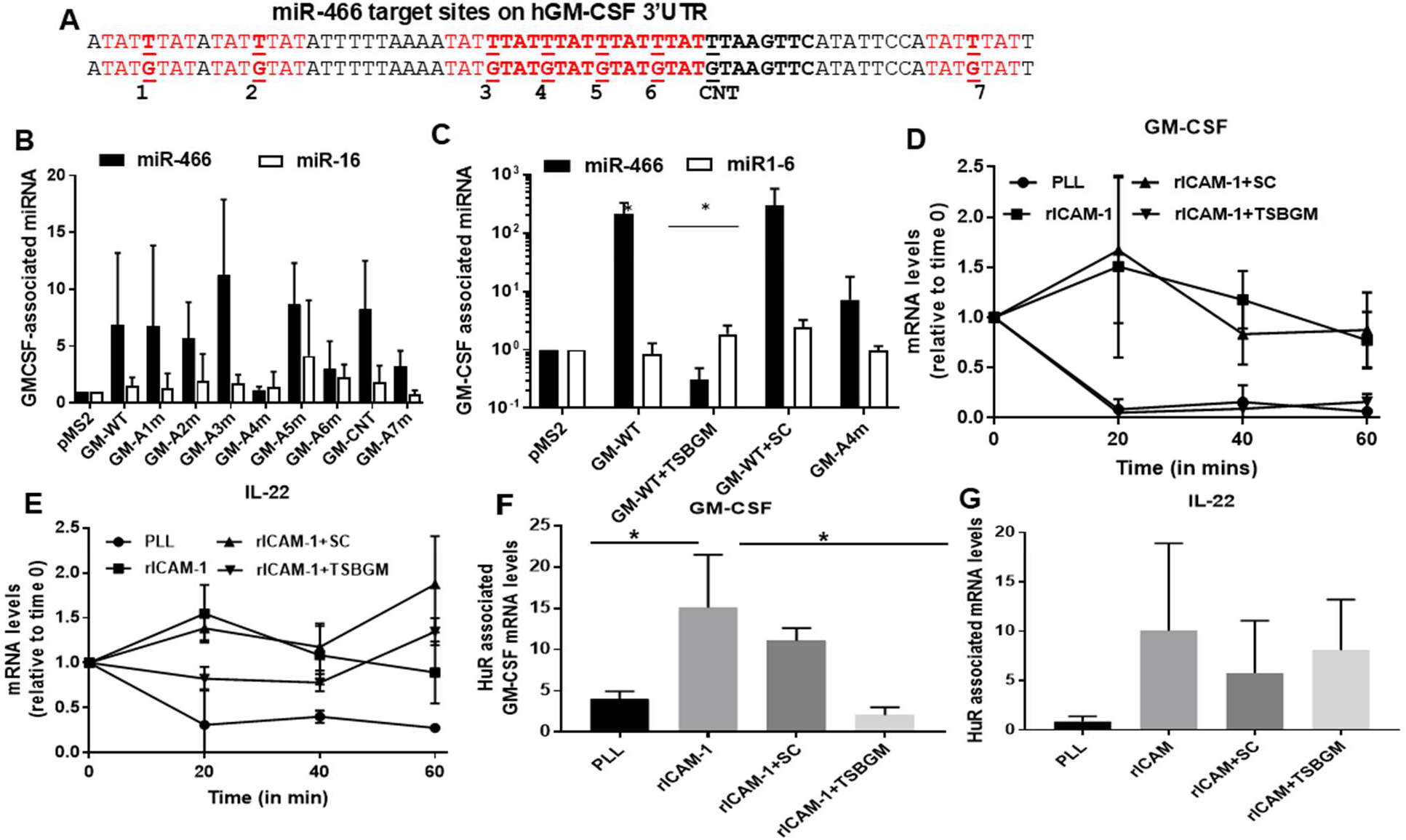
Post-Transcriptional Regulation of GM-CSF by miR-466 and HuR. (***A***) Conserved sequence map of GM-CSF 3-UTR showing the location of site-directed point mutations introduced within the predicted AU-rich elements and CNT (A1m–A7m). (***B***) NIH-3T3 cells were co-transfected with wild-type or A1m–A7m pMS2-GM constructs and pMS2-GST. After 48 hours, cells were crosslinked, lysed, and subjected to GST pull-down. Crosslinks were reversed, RNA was extracted, and miR-466 and miR-16 levels were quantified by qRT-PCR using the MS2-TRAP assay (***C***) TSBGM was designed to specifically block the fourth conserved cARE (A4m) within the GM-CSF 3′UTR. NIH-3T3 cells were co-transfected with pMS2-GM-CSF-WT and either TSBGM or SC oligonucleotide and assessed for miR-466 and miR-16 bound levels as previously described. Statistical significance assessed by two-way ANOVA (*P < 0.005) (**D**) HSB2-T cells were transfected with TSBGM or SC, stimulated with 2 ng/mL PMA on ICAM-1 coated plates, and treated with 0.2 mM DRB to halt transcription. RNA was collected at indicated time points, and GM-CSF and IL-22 mRNA levels were quantified by qRT-PCR, normalized to GAPDH, and plotted relative to time 0. Data represent mean ± S.E.M. from three independent experiments. **(*E,F*)** HSB-2 cells transfected with 100 nM SC or TSBGM were stimulated with 2 ng/mL PMA on PLL- or rICAM-1–coated plates for 2 hours. RNA immunoprecipitation was performed using anti-HuR or IgG1, and bound hGM-CSF, hIL-22, and GAPDH mRNAs were quantified by qRT-PCR. Data represent mean ± SEM from three experiments, normalized to isotype control. Statistical significance assessed by two-way ANOVA (*P < 0.005).

The ability of TSBGM to disrupt miR-466 binding was assessed using the MS2-TRAP assay. 466-3T3 cells were transfected with either the empty pMS2 vector, pMS2-GM-WT alone, or co-transfected with pMS2-GM-WT and 25 nM TSBGM or SC oligonucleotide, as well as with the mutant construct pMS2-GM-cARE4m. In cells transfected with pMS2-GM-WT or pMS2-GM-WT + SC, miR-466 binding was strongly enriched (215–270-fold) relative to the empty vector. Co-transfection with TSBGM significantly reduced miR-466 binding to near-background levels (∼0.5-fold). Similarly, mutation of the fourth conserved ARE (cARE4) reduced miR-466 binding to ∼7.1-fold. Importantly, miR-16 binding was unaffected by TSBGM across all constructs, demonstrating target specificity (Fig. 6D).

To evaluate the functional impact of miR-466 blockade by TSBGM, we performed mRNA stability assays in HSB2-T cells transfected with either 25 nM TSBGM or SC. Following stimulation with 2 ng/mL PMA and LFA-1–mediated adhesion, GM-CSF and IL-22 mRNA levels were measured over time. In cells plated on PLL or transfected with TSBGM and adhered to ICAM-1 coated plates, GM-CSF mRNA underwent rapid decay, with nearly 100% degraded by 20 minutes. In contrast, GM-CSF mRNA was largely stable in untreated and SC-transfected cells on ICAM-1, with only ∼25% decay by 60 minutes. IL-22 mRNA remained stable across all conditions on ICAM-1, confirming the specificity of TSBGM for destabilizing GM-CSF transcripts (Fig. 6E). ELISA of supernatants revealed a dose-dependent decrease in GM-CSF protein with increasing TSBGM concentration (25–100 nM), while SC-treated cells showed no change (Fig. S5).

To determine whether the TSBGM oligonucleotide affects the interaction between HuR and GM-CSF mRNA, we performed HuR-RNA immunoprecipitation (HuR-IP) assays in HSB-2 cells transfected with either TSBGM or a SC control oligonucleotide as previously described. HuR association with GM-CSF mRNA was markedly reduced by TSBGM (∼2 fold) in comparison to untreated cells and SC transfected cells (11-15 fold), indicating that blocking miR-466 binding prevents HuR recruitment (Fig. 6F). HuR binding to IL-22 mRNA was unaffected (Fig. 6G), confirming the selectivity of TSBGM.

Together, these findings extend the cooperative HuR–miR-466 regulatory mechanism to GM-CSF. We identify cARE4 as the critical regulatory hub and demonstrate that TSBGM selectively destabilizes GM-CSF mRNA by blocking miR-466 binding and subsequent HuR association, without disrupting unrelated transcripts.

### miR-466-Mediated Control of IL-23A Expression and Therapeutic Blockade with TSB23

We next investigated whether IL-23A, a key Th17 cytokine, is regulated by miR-466 through a similar mechanism. Notably, overexpression of miR-466 leads to elevated IL-23A mRNA levels in human macrophages^21^. The IL-23A 3′UTR contains four cAREs overlapping predicted miR-466 seed sequences (Fig. 7A). Using the MS2-TRAP system, NIH-3T3 cells were co-transfected with pMS2-IL-23-WT and either aMIR466 or cMIR. miRNA pull-down assays showed a ∼6–8-fold enrichment of miR-466 binding to the IL-23A 3′UTR in cMIR-transfected cells, which was significantly reduced to baseline (∼ 1 fold) by aMIR466, confirming miR-466–specific binding (Fig. 7B). To map the dominant interaction site, we generated four point mutants targeting each of the conserved AREs (A1m–A4m), as well as a non-overlapping control mutant (23-CNT). These mutated pMS-23-WT constructs were transfected into the 466-3T3 stable cell line, and miR-466 binding was assessed. Mutation of the third ARE (PMS-23-A3m) resulted in a substantial reduction (∼4-fold) in miR-466 binding, while mutations in 23-A1m, 23-A2m, and 23-A4m maintained strong enrichment (∼1000-fold) identifying cARE3 as the dominant functional site (Fig. 7C).

**Figure 7.**
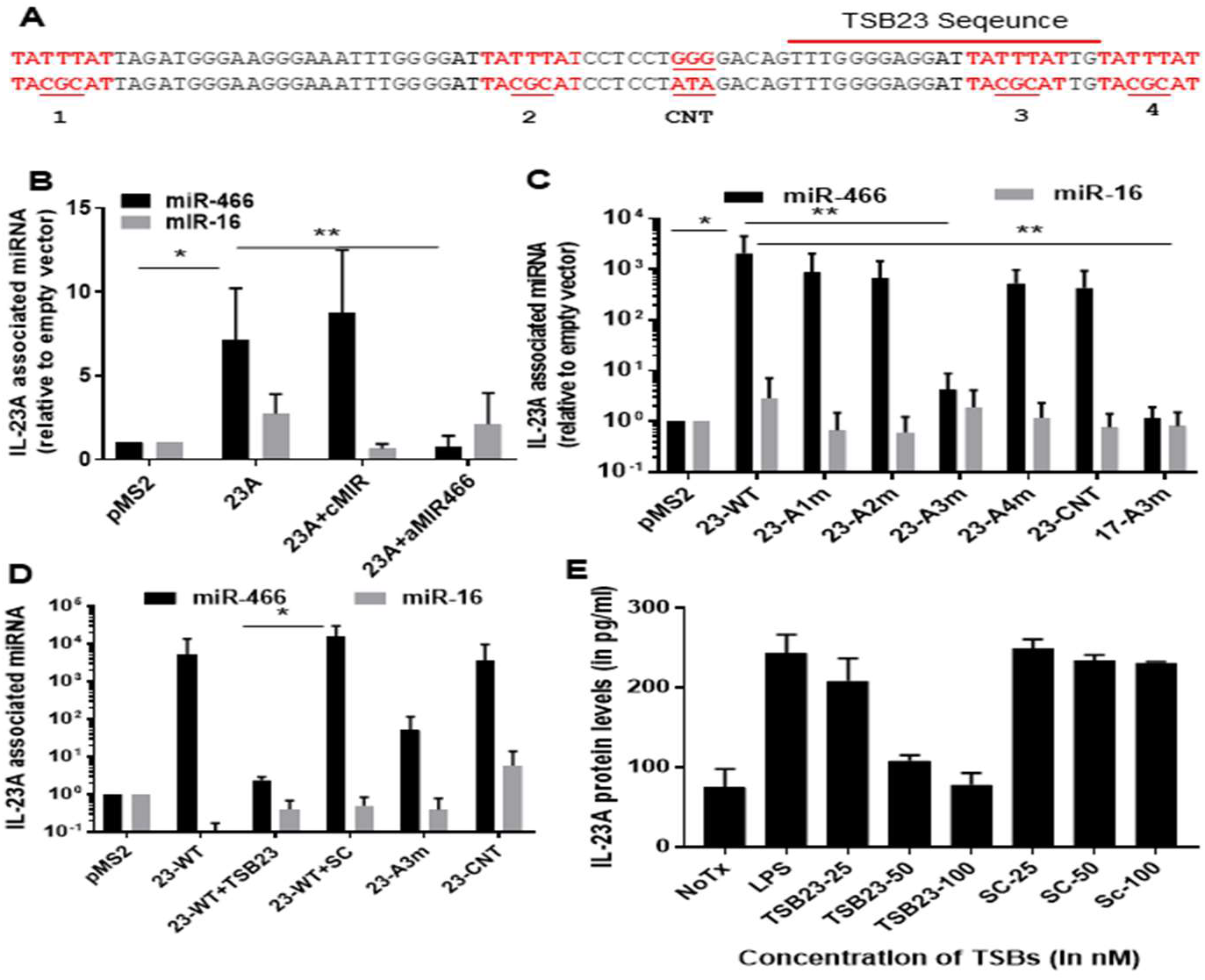
miR-466 specifically binds the IL-23p19 3′UTR at a conserved ARE site and is effectively blocked by TSB23. **(*A*)** Schematic representation of the IL-23p19 3′UTR showing four conserved cAREs. Core ARE motifs and a control mutation site (CNT) are highlighted. ***(B*)**. NIH-3T3 cells were co-transfected with pMS2-23-WT and either 25 nM cMIR or aMIR466. The binding of miR-466 (black bars) and miR-16 (gray bars) was quantified by qRT-PCR as previously described. **(*C*)** Site-directed mutational analysis of each of the four predicted cAREs within the IL-23p19 3′UTR. Constructs harboring individual point mutations (A1m–A4m) were transfected into 466-3T3 cells, and miR-466 binding was quantified via MS2-TRAP and qRT-PCR as previously described. **(*D*)** 466-3T3 cells were co-transfected with pMS2-23-WT and either 25 nM TSB23 or SC. The bound levels of miR-466 and miR-16 was determined by qRT-PCR as previously described. Data were collected from three independent experiments, with all samples analyzed in triplicate. Statistical significance assessed by two-way ANOVA (*P < 0.005) (**P < 0.001)

We designed a TSB (TSB23) targeting the 3^rd^ cARE miR-466 binding site. In MS2-TRAP assays, co-transfection of 466-3T3 cells with pMS2-23-WT and TSB23 reduced miR-466 binding from ∼5100-fold to ∼2.3-fold—greater inhibition than that achieved by mutating cARE3, demonstrating potent and specific blockade (Fig. 7D). miR-466 binding remained high in cells transfected with pMS2-23-WT, pMS2-23-WT+SC, or the pMS2-23-CNT, indicating that TSB23 specifically disrupts the miR-466–cARE3 interaction. Importantly, TSB23 did not affect miR-16 binding to the IL-23 3′UTR, confirming its target specificity. To assess the functional impact of TSB23 on IL-23A protein expression, MHS-2 macrophages were transfected with increasing concentrations (25, 50, and 100 nM) of TSB23 or SC oligonucleotide and stimulated with LPS. IL-23A protein levels in culture supernatants were measured by ELISA 24 hours post-stimulation. LPS stimulation alone markedly increased IL-23A levels to 243 pg/ml compared to 74 pg/ml untreated (NoTx) controls. TSB23 treatment led to a dose-dependent reduction in IL-23A protein, with maximal suppression of 78 pg/ml at 100 nM. In contrast, SC oligonucleotides had no significant effect at any concentration tested (Fig. 7E). These results demonstrate that TSB23 selectively interferes with post-transcriptional regulation of IL-23A, effectively reducing its expression in activated macrophages. These findings identify cARE3 as the primary miR-466 binding site in the IL-23A 3′UTR and demonstrate that TSB23 effectively blocks this interaction, leading to selective suppression of IL-23p19 expression.

### Therapeutic efficacy and cytokine specificity of TSBs targeting IL-17A, IL-23p19, and GM-CSF in MOG₃₅–₅₅/CFA-induced EAE

To assess therapeutic efficacy, we tested TSB23 and TSBGM in the MOG₃₅–₅₅/CFA-induced EAE model in 2D2 TCR transgenice mice, which is less dependent on IL-17A than the zymosan-adjuvanted model. Mice treated with SC oligos developed severe disease, with a peak clinical score of ∼3.6 by day 18. In contrast, TSB23 treatment inhibited disease progression (peak score ∼2), while TSBGM conferred intermediate protection (peak ∼2.5) (Fig. 8A). Importantly, AUC of EAE clinical scores over time confirmed the superiority of IL-23 blockade. TSB23-treated mice exhibited the lowest cumulative disease burden (33.36), followed by TSBGM (43.27), while SC-treated mice showed the highest AUC values (96.44) (Fig. 8B).

**Figure 8.**
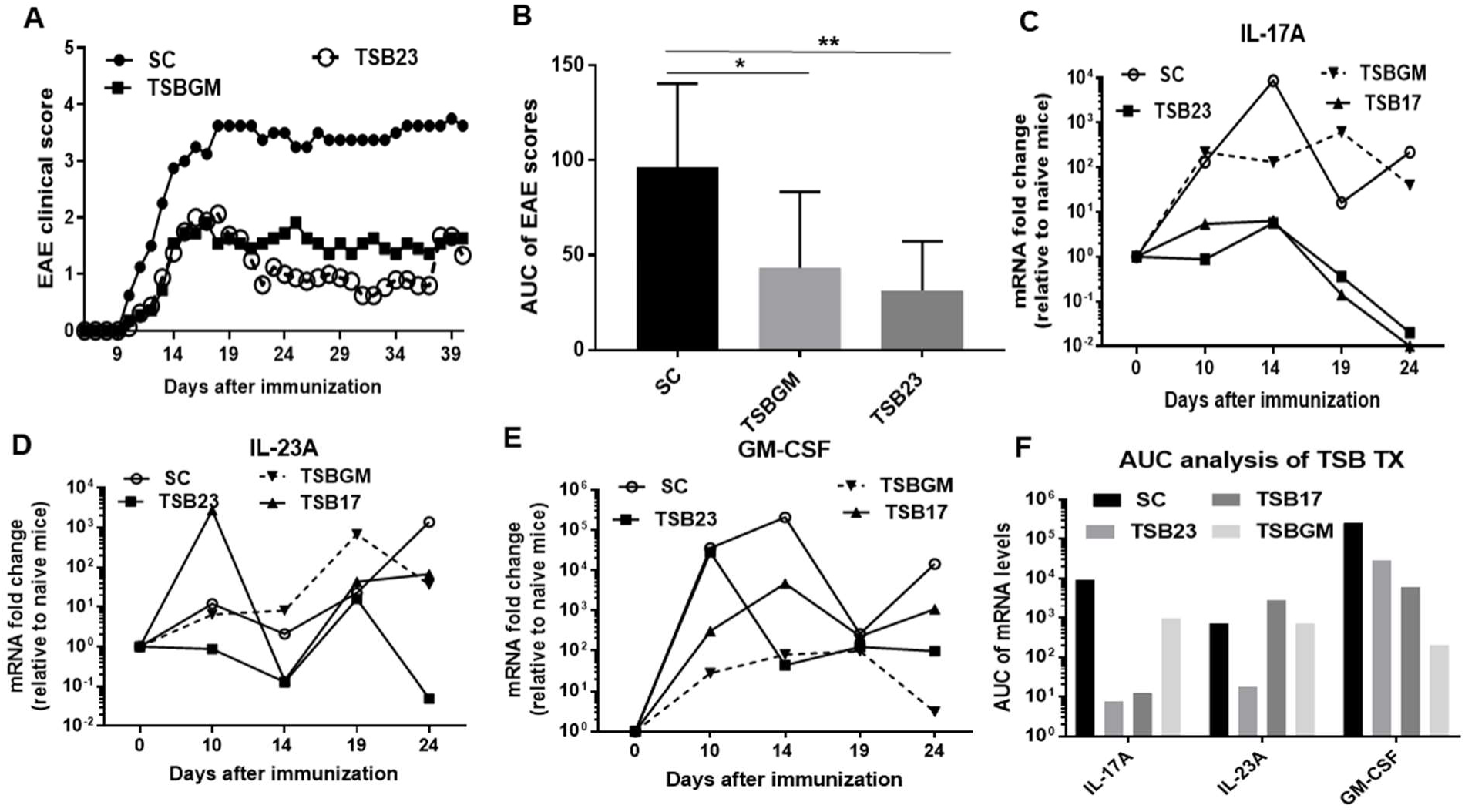
Therapeutic efficacy and cytokine mRNA of TSBs targeting IL-17A, IL-23A, and GM-CSF in MOG₃₅–₅₅/CFA-induced EAE. **(*A*)** EAE was induced in 2D2-transgenic mice via subcutaneous immunization at two sites with an emulsion containing 100 μg of MOG₃₅–₅₅ peptide and 250 μg of CFA per mouse. Treatment with either 5 mg/kg of SC (n= 8), TSB23 (n=16) or TSBGM (n=11) was initiated on day 6 post-immunization. A double dose (10 mg/kg) was administered on day 9, followed by maintenance dosing at 5 mg/kg every three days for the duration of the study. Clinical symptoms were monitored and scored daily. (***B***) Ares under the curve analysis of *A* and with statistical analysis. (***C,D,E***) EAE was induced in 2D2-transgenic mice and treated with TSB17 (n=4), TSB23 (n=4), TSBGM (n=4), or SC oligonucleotide as described above. At days 0, 10, 14, 19, and 24 post-immunization, one mouse per treatment group was sacrificed, and spinal cords were collected for molecular analysis. Total RNA was isolated, reverse-transcribed, and analyzed by qRT-PCR to quantify mRNA levels of IL-17A (***C***), IL-23A (***D***), and GM-CSF (***E***). Transcript levels were normalized to a housekeeping gene and expressed as fold change relative to naïve mice (day 00. Each sample was analyzed in triplicate, and the data are presented for each individual mouse at the corresponding time point. (***F***) AUC analysis of ***C***, ***D*** and ***E***. Statistical significance assessed by two-way ANOVA (*P < 0.005) (**P < 0.001).

We next examined the effect of TSBs on the transcript-levels in the lumbar spinal cords of EAE mice over a time course of 21 days. RNA was collected at days 0, 10, 14, 19, and 24 post-immunization, and cytokine mRNA levels were assessed.

#### IL-17A

In SC-treated animals, IL-17A mRNA levels rose sharply by day 10, peaked at day 14, and gradually declined by day 21. Both TSB17 and TSB23 treatments attenuated IL-17A mRNA expression, particularly at the peak on day 14, indicating that direct targeting (TSB17) and indirect suppression via IL-23 inhibition (TSB23) effectively reduce IL-17A levels (Fig. 8C). This was reflected in the AUC analysis (Fig. 8F), where TSB17 and TSB23 treatments reduced cumulative IL-17A transcript abundance by over an order of magnitude compared to SC controls. TSBGM produced a modest reduction, consistent with limited feedback from GM-CSF on IL-17A regulation (Fig. 8C).

#### IL-23A

TSB23 treatment led to a marked and sustained suppression of IL-23A mRNA levels across all time points, with the most dramatic inhibition observed at day 24. This confirms the specificity and potency of the TSB23 oligonucleotide, as reflected in the AUC, which dropped from ∼10⁴ in SC-treated animals to <10² in TSB23-treated mice. TSB17 and TSBGM showed only minor effects, indicating partial upstream influence. Notably, the secondary IL-23A peak observed in SC-treated mice at day 19 was completely abrogated with TSB23 (Fig. 8D).

#### GM-CSF

TSBGM treatment produced the most pronounced suppression of GM-CSF expression, particularly following its day 10 peak (Fig. 8E). This was captured in the AUC analysis, where GM-CSF levels dropped several logs in TSBGM-treated animals (208) relative to SC (253181). TSB17 and TSB23 also reduced GM-CSF transcript levels to a lesser extent, supporting the notion of cytokine crosstalk within the Th17 network (Fig. 8F).

Together, these findings demonstrate the therapeutic potential of miR-466–targeting TSBs in vivo, with TSB23 showing the strongest efficacy and transcript specificity in the EAE model.

## DISCUSSION

Targeting miRNAs for therapeutic purposes has been hindered by off-target effects and global inhibition of miRNA function. Our study addresses this barrier by introducing a mechanistically informed approach using TSBs to selectively disrupt individual miRNA–mRNA interactions without interfering with global miRNA networks. We identified and therapeutically exploit a previously unrecognized cooperative interaction between miR-466 and the RBP HuR that stabilizes IL-23A/Th17 inflammatory cytokine transcripts. In contrast to the canonical role of miRNAs in mRNA transcript repression, our findings reveal a noncanonical, stabilizing function of miR-466 that can be selectively disrupted to reduce gene expression. This interaction occurs at discrete *cis*-elements that serve as hubs for miRNA-RBP co-regulation. miR-466-interaction with it’s target site motifs within IL-17A, GM-CSF and IL-23A 3′UTRs facilitates HuR binding probably by structural remodeling of the local RNA landscape, creating a favorable binding interface for HuR^38^. Together, these results expand the classical paradigm of miRNA-mediated repression and highlight a novel regulatory axis that can be therapeutically targeted to modulate cytokine expression with precision and specificity.

Our previous work and others’ have established HuR as a post-transcriptional stabilizer of inflammatory transcripts^8,19,39^. Historically, HuR and miRNAs were thought to act antagonistically, competing for overlapping or adjacent binding sites. For example, HuR has been shown to displace miR-200 from ZEB1 and miR-637 from CRP transcripts^10,15^. As a unique miRNA that binds AREs, miR-466 has been shown to influence the expression of key pro-inflammatory cytokines and growth factors—suggesting a critical, yet poorly understood, role in coordinating post-transcriptional gene regulation during inflammatory responses^21^. Here, we identify dominant cAREs within each cytokine transcript—cARE3 in IL-17A, cARE4 in GM-CSF, and cARE3 in IL-23p19 (Fig.4A, 7A & 8A)—as key *cis*-regulatory hubs where miR-466 binding facilitates HuR association. Employing global miR-466 inhibition via antimiRs, transcript-specific occlusion using TSBs, and genetic HuR depletion, we demonstrate that HuR recruitment to target mRNAs is dependent on antecedent miR-466 engagement, as evidenced by a substantial loss of HuR association upon disruption of miR-466–mRNA interactions (Fig. 3E, 4B & 6F). Furthermore, mutation of the IL-17A cARE3 element disrupts both HuR and miR-466–mediated transcript stabilization (Fig. 2E). Importantly, while miR-466 binding to mRNA occurs independently of HuR (Fig.3F), HuR is essential for the stabilizing effect as demonstrated in HuR kno cells (Fig. 1E). These findings contrast with the model proposed by Chen et al., which suggested that HuR stabilizes cytokine transcripts by antagonizing miR-466 activity^40^. Our data reveal that miR-466 facilitates HuR recruitment by directly binding to cAREs, as demonstrated through mutagenesis, RNA pulldown, and MS2-TRAP assays, indicating that miR-466 engagement is a prerequisite for HuR association (Fig. 4B, 6C, 7D).

The miR-466 target sites (UAUUUAU) identified within the 3′UTRs of IL-17A, GM-CSF, and IL-23A serve as key regulatory hubs for miR-466–mediated post-transcriptional control. These U-rich heptamers encompass the canonical AUUUA pentameric motifs traditionally associated with RBPs such as HuR. Although the transcripts examined here harbor both miR-466 target sites and multiple pentameric AUUUA motifs, our data suggest that HuR engagement is not solely dictated by the presence of these canonical pentamers. For instance, Bcl-2 mRNA contains several AUUUA motifs but only a single miR-466 target site (UAUUUAU); targeted blockade of this site using a Bcl-2 TSB abrogated HuR binding and led to reduced mRNA levels (unpublished data). This implies that HuR does not inherently bind to all AUUUA motifs, and that occupancy of UAUUUAU by miR-466 may be necessary to remodel local RNA secondary structure or recruit auxiliary factors that facilitate HuR loading. These findings challenge the classical view that AUUUA motifs are sufficient for HuR binding and support a revised model in which miR-466 binding to composite ARE motifs primes the transcript for HuR engagement.

Previous studies have established HuR as a stabilizing RBP that counteracts miRISC-mediated repression. For instance, Kundu et al. demonstrated that HuR promotes the stability of inflammatory transcripts such as SphK1 by binding to AREs and displacing AGO2–miRNA complexes from the 3′UTR, supporting a classical antagonistic model in which HuR and miRISC act in opposition—HuR protecting transcripts from decay, and AGO2-bound miRNAs promoting degradation^41^. Similar antagonism has been reported across multiple cancers, where HuR competes with miRNAs such as miR-16, miR-122, or miR-331-3p to inhibit miRISC function and facilitate mRNA translation^42^. Conversely, HuR can also facilitate AGO2-mediated repression. In cervical cancer cells, HuR promotes *let-7* loading onto the *MYC* 3′UTR, enhancing silencing^15^. In neurons, HuR synergizes with miR-26/AGO2 to suppress *Rgs4*, a gene implicated in tumorigenesis^43^. Further complexity arises from recent transcriptome-wide analyses by Li et al., which show that ∼10% of AGO2 sites overlap with ∼18% of HuR-bound regions. Such co-occurrence supports a model of combinatorial regulation, where HuR may either block or enhance miRNA/AGO2 function depending on *cis*-regulatory context. For example, HuR antagonizes miRNA repression of *BTG2* and *CDK16*, but cooperates with miRISC to regulate *MSMO1* expression^44^. Importantly, knockdown of either protein led to decreased stability of *MSMO1*, suggesting a cooperative mechanism in some settings. Indeed, AGO2 itself may persist on transcripts in a non-degradative state, potentially promoting changes in the mRNA folding structure that facilitate HuR or other stabilizing RBPs binding. Our findings build on this cooperative model by demonstrating that miR-466, most likely via AGO2-loaded miRISC, facilitates HuR recruitment to cytokine mRNAs, including *IL-17A* and *GM-CSF*. While miR-466 binding to its target site occurs independently of HuR, HuR association is abolished by either miR-466 neutralization or TSB-mediated blockade of the seed site. These data support a model in which miRISC engagement creates a structural or kinetic environment that facilitates HuR binding, thereby promoting mRNA stabilization. By targeting the miR-466 target site, TSBs disrupt this cooperative interaction, preventing miRISC-mRNA interaction and HuR recruitment. This represents the mechanism of action for TSBs—not merely as inhibitors of miRNA-mediated stabilization, but as precision tools to dismantle pro-stabilizing RNA–protein complexes on selected transcripts.

A major challenge in oligonucleotide-based therapeutics is minimizing off-target effects, particularly when targeting broadly expressed miRNAs like miR-466. TSB treatment selectively reduced target mRNA levels with minimal impact on unrelated transcripts. TSB23 significantly suppressed IL-23A mRNA and, consistent with its upstream role, also reduced IL-17A and GM-CSF expression (Fig.8F). TSB17 decreased IL-17A without altering IL-23A levels, while modest downregulation of GM-CSF. We observed (Fig. S1E-F) and previously proposed that IL-17A expression precedes GM-CSF induction in the MOG₃₅–₅₅ EAE model, likely accounting for the observed cytokine interdependence ^45^. TSBGM treatment resulted in robust GM-CSF silencing with no effect on IL-23A and limited changes in IL-17A, again suggesting downstream effects rather than direct targeting (Fig. 8F). Mechanistically, MS2-TRAP assays demonstrated that TSB17, TSB23, and TSBGM each blocked miR-466 binding to their respective targets without interfering with miR-16—a miRNA that also targets AREs—thus excluding nonspecific displacement of miRISC complexes (Figs. 4B, 6C, 7D). This specificity was further validated functionally: TSBGM suppressed GM-CSF but not IL-22, and TSB17 downregulated IL-17A but not IL-6 (Fig. 6E & S4B). Previous studies have demonstrated the target specificity of a range of TSBs through RNA-sequence analysis, functional assays, and downstream validation. For examples. TSB targeting the miR-155 site on Arg2 restored Arg2 expression and reduced macrophage-driven inflammation^46^; in melanoma, blockade of the miR-16 site on TYRP1 relieved translational repression and limited tumor growth^47^; and in cystic fibrosis, TSBs against CFTR 3′UTR sites reversed miRNA-mediated suppression and rescued protein function^48^. Collectively, these data reinforce the utility of TSBs in selectively disrupting disease-relevant interactions without broadly antagonizing the miRNA or other miRNA-targeted transcripts, offering a therapeutic advantage over conventional anti-mirs.

In oligonucleotide therapeutics, phosphorothioate (PS)-modified backbones involve the substitution of a non-bridging oxygen in the phosphate group with a sulfur atom, which improves nuclease resistance and increases binding to plasma proteins. This modification enhances both serum stability and cellular uptake. In our murine EAE models, the systemic efficacy of TSBs was made possible by incorporating a PS backbone. For translational use, PS-TSBs formulated in lipid nanoparticles or complexed with cell-penetrating peptides offer favorable pharmacokinetics and reduced innate immune activation. These delivery modalities will be critical for human applications where tissue penetration and biodistribution dictate therapeutic efficacy. Notably, the specificity and efficacy of TSBs extended beyond murine systems. Equivalent suppression of IL-17A and GM-CSF was observed in activated primary human T cells transfected with species-specific TSBs (Fig. 4C and 6D), suggesting that the miR-466/HuR regulatory axis is conserved across species and underscoring the translational relevance of this strategy. Supporting this, independent functional studies have demonstrated that transfection of miR-466 mimics or anti-mir-466 oligonucleotides in human cells elicits measurable regulatory effects, further confirming the functional relevance of miR-466 in human post-transcriptional gene regulation^20–22,49^.

Antisense oligonucleotides (ASOs) are synthetic strands of nucleic acids that bind to target mRNAs through Watson–Crick base pairing. In gapmer configurations, a central DNA core flanked by chemically modified ribonucleotides enables recruitment of RNase H1, which cleaves the RNA strand and promotes transcript degradation^50^. ASO efficacy depends on factors such as target site accessibility, duplex stability, and cellular levels of RNase H1⁵⁰. TSBs designed in a similar gapmer format also harnesses RNase H1 to mediate mRNA destabilization. In RNase H1 inhibition experiments, we observed that TSB-induced decay of Bcl2 mRNA was abrogated, indicating that the mechanism of action involves RNase H1–dependent cleavage (data not shown).

A major advantage of TSBs over ASOs or RNAi lies in their design precision. According to the n-Lorem Foundation’s ASO discovery pipeline of 400-500, this initial pool undergoes multiple rounds of *in vitro* testing for potency and specificity, followed by in vivo screening for tolerability and immunogenicity. Even with sophisticated design tools, this multi-step process typically yields only 20–75 candidates that are suitable for preclinical testing, reflecting the considerable time, cost, and optimization burden associated with ASO development. For example, in the case of IL-23A, we identified the preferred miR-466 target site through site-directed mutagenesis and designed TSB23 oligonucleotide to block this interaction—without relying on mRNA stabilization assays or HuR immunoprecipitation experiments. Despite the absence of extensive biochemical validation at the outset, the TSB effectively downregulated IL-23 expression, validating the predictive design approach. We have since simplified this process further, demonstrating that TSBs designed solely against in silico–predicted miR-466 target sites can robustly suppress the expression of several genes, including pro-inflammatory cytokines such as IL-6, TNF-α, and IL-22, as well as other key regulatory targets like *BCL-2*, *NOTCH1*, and *MECP2* (data not shown). In most cases, only a small number of candidate oligonucleotides are needed per target, thereby reducing complexity, enhancing precision, and accelerating therapeutic advancement. Only a small subset of miRNAs function as mRNA stabilizers, while the vast majority promote transcript destabilization. As a result, identifying and inhibiting stabilizing miRNAs by TSB is relatively straightforward. In contrast, when designing TSBs to enhance mRNA stability, multiple destabilizing miRNAs may target the same transcript, making it more challenging to pinpoint a single dominant miRNA to block effectively. This underscores a key advantage of TSBs over traditional ASOs: they act on defined miRNA–mRNA interactions, enabling a focused and rational design strategy.

Despite its central role in regulating mRNA stability, translation, and localization, HuR has long been considered an undruggable target due to its ubiquitous expression, structural redundancy, and involvement in essential cellular processes such as cell cycle progression, DNA repair, and stress response. Direct inhibition of HuR risks widespread toxicity and disruption of homeostatic gene expression. By using a TSB to occlude the miR-466 binding site, we indirectly prevent HuR recruitment to inflammatory transcripts such as IL-17A and GM-CSF, leading to selective mRNA destabilization. This strategy avoids direct inhibition of HuR protein itself and instead leverages transcript-specific context to disarm HuR function in a highly targeted manner. Thus, TSBs offer a precision tool to neutralize HuR activity on pathogenic mRNAs.

Although our target-level validation confirmed selective modulation of IL-17A, GM-CSF, and IL-23A mRNAs, a key limitation of this study is the absence of transcriptome-wide analyses to fully assess off-target effects and validate global precision. Given the broad target landscape of miR-466, future studies incorporating RNA-seq or CLIP-seq will be essential to identify potential context-dependent transcriptomic changes in diverse immune and tissue settings. Additionally, the lack of AGO2-RIP or CLIP experiments limits our ability to biochemically confirm the involvement of RISC components in the observed mRNA stabilization phenotype. These mechanistic and transcriptome-wide assessments will be critical for advancing TSBs as selective, mechanism-informed therapeutics capable of disrupting disease-relevant miRNA–mRNA interactions without globally impairing miRNA function.

## MATERIALS AND METHODS

### Human peripheral T-cell isolation

Peripheral blood samples were obtained from healthy adult donors after written informed consent under protocols approved by the Yale University Institutional Review Board. Peripheral blood mononuclear cells (PBMCs) were isolated by density-gradient centrifugation using Ficoll-Histopaque 1077 (Sigma-Aldrich, St. Louis, MO) CD3⁺ T cells were purified by negative selection using a human T-cell isolation kit (Miltenyi Biotec, Auburn, CA) according to the manufacturer’s instructions. Purity was routinely >95% as assessed by flow cytometry using anti-CD3 antibodies. T cells were activated with anti-CD3 monoclonal antibody (OKT3, 1 µg/mL) for 72 h before use in experiments.

All experiments using human cells were performed with cells from at least three independent donors.

### Cell lines and culture conditions

Jurkat, EL-4, NIH-3T3, MHS-2, and HSB-2 cell lines were obtained from in-house stocks and maintained under standard conditions at 37 °C in a humidified incubator with 5% CO₂. Jurkat, EL-4, HSB-2, and MHS-2 cells were cultured in RPMI-1640 medium (Gibco) supplemented with10% fetal bovine serum (FBS),2 mM L-glutamine,100 U/mL penicillin,100 µg/mL streptomycin.NIH-3T3 cells were cultured in DMEM (Gibco) supplemented with 10% FBS,2 mM L-glutamine, penicillin/streptomycin. Stable transfectants expressing miR-466l-3p (466-3T3, 466-EL-4) or control constructs were generated by plasmid transfection followed by puromycin selection (2–7 µg/mL) for ≥2 week

### Transient transfection and generation of stable cell lines

miR-466l-3p antimiRs (aMIR466) and control antimiRs (aMIR) were obtained from Exiqon (Denmark). Human, mouse T or Jurkat cells were transfected via electroporation using AMAXA nucleofection kits specific for each cell type, following the manufacturer’s protocol (Lonza, Walkersville, MD). Primary T cells were transfected with 25 nM or as indicated concentration of each oligonucleotide. Jurkat cells were transfected with 5 ug of pBBB-17WT or pBBB-17cARE3m and 2.5 ug peGFP. To generate stable EL-4 cell transfectants, cells were electroporated with 2 µg of plasmid DNA using the AMAXA kit (Lonza) and subsequently selected in medium containing 7 µg/ml puromycin for two weeks. Stable NIH 3T3 transfectants were generated by transfecting cells with 2 µg of plasmid DNA using Lipofectamine (ThermoFisher Scientific), followed by selection with 2 µg/ml puromycin. All stable transfectants were maintained in puromycin-containing media for several weeks to ensure sustained selection.

### RNA Immunoprecipitation (RIP) Assay

Primary human T cells (1 × 10⁷ cells per condition) were stimulated with 2 ng/ml PMA for 2 hours on plates coated with either recombinant human ICAM-1 (rhICAM-1) or poly-L-lysine (PLL). Cells were then collected, washed, and resuspended in calcium- and magnesium-free PBS (CMF-PBS). Reversible crosslinking was performed using 1% formaldehyde for 10 minutes at room temperature. Following crosslinking, cells were lysed by sonication at 70% amplitude using a Sonic Dismembrator 500 (Fisher Scientific, Pittsburgh, PA) in immunoprecipitation buffer (25 mM HEPES, pH 8.0; 150 mM KCl; 2.5 mM EDTA; 840 µg/ml NaF; 1 mM DTT; 0.1% NP-40) supplemented with EDTA-free protease inhibitors and 2 U/µl RNasin ((Promega, Madison, WI). Lysates were immunoprecipitated overnight at 4°C with either isotype control (IgG1) or anti-HuR antibodies pre-conjugated to protein A/G Plus agarose beads. Beads were subsequently washed with PBS, and immunoprecipitates were treated with 30 µg/ml proteinase K for 60 minutes at 55°C to reverse crosslinking and digest proteins. RNA was isolated using the Zymo RNA Isolation Kit (Zymo Research, Irvine, CA), reverse transcribed, and analyzed by qPCR in triplicates to assess IL-17A and GAPDH mRNA levels. IL-17A mRNA abundance in HuR immunoprecipitates was normalized to GAPDH. Results are shown as fold enrichment relative to IgG control.

### LFA-1 mediated mRNA stabilization experiments

Petri dishes were coated with either human or mouse recombinant ICAM-1 (rICAM-1) as previously described^9^. For control conditions, dishes were coated with 20 µg/ml poly-L-lysine (PLL) overnight at 4°C. In both human and mouse primary T cells and transfected Jurkat cells, gene transcription and LFA-1 activation were triggered by adding 2 ng/ml PMA for 2 hours on either rICAM-1 or PLL-coated surfaces. EL-4 cells were resuspended at 10⁷ cells/ml in LFA-1 activation buffer (100 mM Tris-HCl, pH 7.5, 0.9% NaCl, 2 mM MnCl₂, 2 mM MgCl₂, 5 mM D-glucose, 1.5% BSA) and incubated on immobilized rICAM-1 for 30 minutes. Following adhesion, the activation buffer was aspirated and replaced with RPMI medium supplemented with 10% FBS. Transcription was inhibited at time zero (T = 0) by the addition of 2 mM DRB (5,6-dichloro-1-β-D-ribofuranosylbenzimidazole) (Sigma-Aldrich, St. Louis, MO), and total RNA was harvested at the indicated time points. Reverse transcription was performed using the iSCRIPT cDNA synthesis kit (Bio-Rad,Hercules, CA). Transcript levels were quantified by quantitative PCR (qPCR) using GAPDH as a normalization control. Relative mRNA stability was calculated using the 2(-ΔΔCt) method, with T = 0 serving as the reference time point.

### Mouse strains, naive T cell isolation and Th17 cell differentiation

C57BL/6 mice were purchased from the Jackson Laboratory (Bar Harbor, ME). RORγt-Cre mice were a generous gift from Dr. Flavell with permission from Dr. Littman^34,51^. The HuR^fl/fl^ were developed in our lab and have been described before^19^. The HuR^fl/fl^ and RORγt-Cre mouse strains were crossed. The RORgt-Cre+HuR^fl/fl^ mice developed normally with 90-95% peripheral T cells lacking HuR.

### Experimental Autoimmune Encephalomyelitis (EAE) Induction and TSB Treatment

EAE was induced in 8–10-week-old mice that were either purchased (Jackson Laboratory) or bred in our mouse facility. A standard MOG₃₅–₅₅ peptide immunization protocol was used to induce. Briefly, mice were injected subcutaneously at two sites over the flanks with 250 μg of MOG₃₅–₅₅ peptide (MEVGWYRSPFSRVVHLYRNGK, GenScript (Piscataway, NJ) emulsified in Complete Freund’s Adjuvant (CFA; Sigma-Aldrich (St. Louis, MO), containing 4 mg/mL heat-killed Mycobacterium tuberculosis H37Ra) or 500 ug Zymosan^52^. Additionally, 200 ng of pertussis toxin (List Biological Laboratories, Campbell, CA) was administered intraperitoneally on days 0 and 2 post-immunization. Mice were monitored daily and scored for clinical signs of disease using a 0–5 scale: 0 = no signs; 1 = complete tail paralysis; 2 = Moderate hind limb weakness (wobbly gait); 3 = Complete hind limb paralysis ;4 = Severe forelimb weakness or paralysis;5 = Minimal movement, severe weakness.

TSB oligonucleotides were synthesized as fully modified phosphorothioate oligos with LNA (Qiagen). Mice were randomly assigned to receive either SC or TSBs targeting IL-17A, IL-23p19, or GM-CSF. TSBs were administered by intraperitoneal injection at a dose of 5 mg/kg in PBS, beginning on day 6 post-immunization (pre-onset) and a double dose on day 9 (10 mg/kg), and repeated every 3 days with a single dose (5 mg/kg) thereafter till the end of the experiment. Control animals received equal volumes of PBS or scrambled control oligo.

Clinical scores were recorded daily, and mice were euthanized at peak disease or as otherwise specified for tissue collection. For cytokine expression analysis, lumbar spinal cords were harvested, snap-frozen in liquid nitrogen, and processed for RNA extraction using TRIzol (Thermo Fisher Scientific, Waltham, MA). Cytokine mRNA levels were quantified by qRT-PCR using SYBR Green chemistry (Bio-Rad) and normalized to GAPDH. Fold induction was calculated using the 2–ΔΔCt method relative to naïve controls. For samples with undetectable Ct values, a cycle threshold of 50 was assigned, corresponding to the total number of amplification cycles. TSB specificity and efficacy were determined by assessing reductions in the expression of the intended target cytokine without off-target suppression of unrelated transcripts. All animal procedures were approved by the Yale IACUC and carried out in accordance with NIH guidelines.

### Ex Vivo Expansion of MOG-Specific T Cells

MOG-specific splenic T cells were expanded *ex vivo* derived from immunized mice. Briefly, mice were euthanized at day 21, peak of EAE scores. Spleens were harvested, mechanically dissociated through a 70 µm cell strainer, and red blood cells were lysed using ACK lysis buffer (Thermo Fisher Scientific). Single-cell suspensions were washed and resuspended in complete RPMI-1640 medium supplemented with 10% fetal bovine serum (FBS), 2 mM L-glutamine, 1 mM sodium pyruvate, 0.1 mM non-essential amino acids, 50 µM β-mercaptoethanol, and 100 U/mL penicillin-streptomycin. Cells were plated at 2 × 10⁶ cells/mL in 24-well plates and pulsed with 10 µg/mL MOG₃₅–₅₅ peptide (GenScript). Cultures were maintained at 37°C with 5% CO₂ for 3–5 days. On day 3 or 5, cells were harvested, washed, and used for downstream assays.

### Intracellular IL-17A Staining in 2D2 Primary T Cells and EL-4 Cells

T cells were isolated from spleens of MOG₃₅–₅₅–immunized 2D2 transgenic mice at peak EAE (day 21 post-immunization) and cultured in RPMI-1640 supplemented with 10% fetal bovine serum (FBS), 2 mM L-glutamine, and antibiotics. Cells were stimulated with 10 µg/mL MOG₃₅–₅₅ peptide (GenScript) for 3–5 days to enrich for antigen-specific populations. Cells were transfected with 25 TSB by electroporation using the cell specific Lonza kit. Cells were stimulated with 2 ng/mL PMA and seeded onto ICAM-1–coated plates. After 6 hours of stimulation in the presence of 5 μg/mL Brefeldin A (BioLegend), cells were fixed with 4% paraformaldehyde for intracellular cytokine staining. Permeabilization and staining were performed using the BD Cytofix/Cytoperm™ kit BD B (San Jose, CA) with anti-mouse IL-17A–PE antibody (BioLegend, clone TC11-18H10.1).

EL-4 murine T lymphoma cells were maintained under the same culture conditions. For cytokine induction, 1 × 10⁶ cells/mL were stimulated with 50 ng/mL PMA and 1 μg/mL ionomycin (Sigma-Aldrich) for 6 hours in the presence of 5 μg/mL Brefeldin A. After stimulation, cells were washed, stained with Fixable Viability Dye eFluor 506 (Thermo Fisher) to exclude dead cells, and processed for intracellular IL-17A staining as described above.

Samples were analyzed on a BD LSRFortessa™ flow cytometer, and data were processed using FlowJo™ software (BD Biosciences). Results represent one of three independent experiments and are reported as either percentage of IL-17A⁺ cells or geometric mean fluorescence intensity (gMFI).

### Plasmids, cloning and reporter constructs

The pBBB, pEGFP, pLMP and pMIRGLO and plasmids were available in our lab. The pMS2 and pMS2-GST plasmids were a kind gift of Dr. Myriam Gorospe^36^. The human IL-17A 3’-UTR was obtained by PCR of cDNA derived from stimulated human T cells. The IL-17A 3’-UTR was cloned using the INFUSION kit from Clontech (Mountain View, CA) in the pBBB plasmid after linearization with the BamH1 restriction enzyme. The ARE and NARE sequences were generated by PCR from the cloned IL-17A 3’-UTR in pBBB. These fragments were also cloned in the pBBB as described above. The pMS2 plasmid was linearized with the EcoR1 restriction enzyme. The IL-17A 3’-UTR fragments were cloned in the pMS2 plasmids using the INFUSION kit as described above. The miR-466l-3p gene was generated by PCR from cDNA of mouse T cells. These were cloned in the pLMP plasmids that was linearized with Hpa1 and EcoR1 restriction enzymes.

The 3′ untranslated regions (3′UTRs) of human IL-23p19 (Il23a) and GM-CSF (Csf2) were amplified from total RNA extracted from PBMCs tissues using reverse transcription followed by PCR. cDNA synthesis was performed using SuperScript III Reverse Transcriptase (Invitrogen), and gene-specific primers were designed to amplify the full-length 3′UTRs based. PCR products were cloned into the pBBB vector downstream of the β-globin reporter using XhoI and NotI restriction sites. Ligation reactions were performed using T4 DNA Ligase (New England Biolabs), and constructs were transformed into *E. coli* DH5α competent cells. Positive colonies were screened by colony PCR and verified by Sanger sequencing. Site-directed mutagenesis of cAREs within the 3′UTRs was performed using the QuikChange II XL Site-Directed Mutagenesis Kit (Agilent Technologies, Santa Clara, CA) according to the manufacturer’s protocol. All constructs were sequence-verified prior to transfection into NIH 3T3 or EL-4 cells for downstream functional assays.

### Site-Directed Mutagenesis of AREs and Control Sequences

Individual cARE motifs were disrupted using site-directed mutagenesis. Point mutations were introduced to convert the canonical AU**U**UA motif to a non-functional variant (AU**G**UA). Mutagenesis was performed using a high-fidelity DNA polymerase (Agilent Technologies, CA, USA) with custom-designed primers harboring the desired nucleotide substitutions. Following PCR amplification, reaction products were digested with DpnI (New England Biolabs) to eliminate methylated parental plasmid templates. The resulting plasmids were transformed into chemically competent *E. coli* cells for propagation. Colonies were screened, and plasmids were purified and verifie d by Sanger sequencing to confirm the presence and precision of the intended mutations.

### mRNA Decay Assay Using β-Globin Reporter Constructs

NIH 3T3 cells were seeded at a density of 0.5 × 10⁶ cells/ml in 12-well plates. Cells were co-transfected with 2 µg of pBBB or its variant plasmids and 1 µg of pEGFP plasmid using Lipofectamine (Invitrogen), according to the manufacturer’s protocol. Following transfection, cells were serum-starved for 24 hours, then stimulated with medium containing 20% fetal bovine serum (FBS) for 1 hour. Total RNA was harvested at hourly intervals up to 4 hours post-stimulation. Reverse transcription was performed as previously described, and mRNA levels were quantified by qPCR. β-globin mRNA levels were normalized to eGFP mRNA to control for transfection efficiency. Fold changes in β-globin expression were calculated relative to the 1-hour time point following serum stimulation.

### miRNA Pulldown Assay Using pMS2-Tagged Constructs

Cells were co-transfected with plasmids encoding either the pMS2 vector containing the 3′UTR of interest or the empty pMS2 vector, along with pMS2-GST at a mass ratio of 2:1 (i.e., two-thirds pMS2 + 3′UTR or pMS2 alone, and one-third pMS2-GST). Transfections were performed using Lipofectamine, electroporation or specify reagent following the manufacturer’s instructions, and cells were incubated overnight to allow expression of the tagged transcripts and the MS2-GST fusion protein. The following day, cells were subjected to reversible crosslinking using 1% formaldehyde for 10 minutes at room temperature to preserve RNA-protein interactions. Crosslinking was quenched with glycine (final concentration 125 mM), and cells were washed with cold PBS and collected by centrifugation. Cell pellets were lysed by sonication at 70% amplitude using a Sonic Dismembrator 500 (Fisher Scientific, Pittsburgh, PA) in ice-cold immunoprecipitation buffer (25 mM HEPES, pH 8.0; 150 mM KCl; 2.5 mM EDTA; 840 µg/ml NaF; 1 mM DTT; 0.1% NP-40) supplemented with EDTA-free protease inhibitors and 2 U/µl RNasin (Promega). Lysates were clarified by centrifugation at 14,000 × g for 15 minutes at 4°C to remove cellular debris. The resulting supernatants, containing RNA-protein complexes, were incubated with glutathione-conjugated Sepharose 4B beads (GE Healthcare) pre-equilibrated in immunoprecipitation buffer. The pulldown reactions were rotated overnight at 4°C to allow specific binding of MS2-GST fusion proteins and their associated mRNAs to the glutathione beads. Beads were then washed five times with high-salt buffer (immunoprecipitation buffer containing 300 mM KCl) to reduce nonspecific interactions. After the final wash, miRNA was eluted by proteinase K digestion (30 µg/ml in elution buffer) for 45 minutes at 55°C to reverse the crosslinking and release miRNA from the complexes. miRNA was subsequently isolated using the Qiagens, miREasy kit according to the manufacturer’s instructions. miRNA was reverse transcribed using the Qiagen QuantiTect Reverse Transcription Kit and subjected to reverse transcription and quantitative PCR (qPCR) analysis to detect specific miRNAs enriched in the pulldown.

### Dual-Glo Luciferase Reporter Assay

The human IL-17A 3’-UTR was cloned in the Sac1 restriction site of the pMIRGLO dual luciferase vector ((Promega, Madison, WI, USA). The 17A-MTA1, 17A-MTA2 and 17A-MTA3 mutants of the human IL-17A 3’-UTR were created by site directed mutagenesis as described earlier. NIH 3T3 cells were cultured overnight at a concentration of 5.10^5^ cells/ml in a 12 well plate. Cells were transfected with the pMIR, pMIR-h17A, pMIR-MTA1, pMIR-MTA2 and pMIR-MTA3 plasmids using Fugene HD reagent (Promega) and left overnight. The cells were stimulated with 1ug/ml Anisomycin for 6 hours, after which cells were harvested and analyzed for Firefly luciferase and Renilla luciferase according to the manufacturer’s instructions. The Firefly luciferase levels were normalized relative to Renillia luciferase levels. Results represent mean of ≥3 experiments.

### Imiquimod-Induced Psoriasis Model and Topical TSB Treatment

Mice were shaved approximately 2 cm × 2 cm 48 hours prior to initiation of treatment. Psoriasiform skin inflammation was induced in 8–10-week-old male and female mice by daily topical application of 5% imiquimod (IMQ) cream (Aldara™, 3M Pharmaceuticals, St. Paul, MN) to the shaved dorsal skin, following established protocols. Mice received a daily dose of 62.5 mg IMQ cream applied uniformly to the shaved back skin and 10 mg to each ear for 5 consecutive days.

Animals were randomly assigned to treatment groups receiving one of the following topical formulations:

- SC Oligonucleotides
- TSB17

TSB oligonucleotides were synthesized as fully phosphorothioated and 2′-O-methyl–modified oligonucleotides incorporating locked nucleic acids (LNAs) by Qiagen. Oligonucleotides were formulated in 10% (w/v) Pluronic F-127 dissolved in PBS, a thermosensitive hydrogel that remains liquid at cold temperatures and gels upon application to the skin at body temperature. A total of 50 µg n 50 µl of TSB formulation or SC oligonucleotides were applied topically 30 minutes after IMQ application once daily, for a total of five treatments. Clinical severity was evaluated daily using a modified PASI (Psoriasis Area and Severity Index) scoring system assessing erythema and scaling on 0–4 scale. At day 6, mice were euthanized and dorsal skin and ear tissues were harvested for RNA protein analysis. Cytokine expression (IL-17A, IL-23A, GM-CSF) was quantified by qRT-PCR, normalized to GAPDH, and expressed relative to SC controls.

All animal procedures were approved by the Yale IACUC and conducted in compliance with NIH guidelines for the care and use of laboratory animals.

### TSB and Anti-miR Oligonucleotides

TSBs and SC oligonucleotides were synthesized as fully phosphorothioate-modified antisense oligonucleotides incorporating locked nucleic acid (LNA) residues to enhance target affinity and nuclease resistance. All TSB sequences were designed to overlap the miR-466 seed-binding regions within the 3′UTRs of specific target transcripts and were synthesized and HPLC-purified by Qiagen (Hilden, Germany). Lyophilized oligonucleotides were reconstituted in nuclease-free water to a stock concentration of 100 µM and stored at −20°C until use.

Antisense oligonucleotide targeting miR-466l-3p (antiMIR-466); Cat. No. AM17000, Ambion, Thermo Fisher Scientific). A scrambled anti-miR negative control with no known mammalian targets (cMIR; Cat. No. AM17010, Ambion, Thermo Fisher Scientific).) was used as a specificity control. Both oligonucleotides were resuspended in nuclease-free water to a final stock concentration of 100 µM. Cells were transfected with 25–100 nM of either aMIR466 or cMIR using FuGENE HD Transfection Reagent (Promega) according to the manufacturer’s instructions. Transfection efficiency and miRNA inhibition were confirmed by qRT-PCR and functional readouts. Treated cells were subsequently used in downstream assays including mRNA decay, RNA immunoprecipitation (HuR-RIP), and cytokine expression analysis

### LPS inflammation model

C57BL/6 wild-type and IL-17A⁻/⁻ mice (8–10 weeks old, male and female) were housed under specific pathogen-free conditions with ad libitum access to food and water. All animal procedures were conducted in accordance with protocols approved by the Yale University Institutional Animal Care and Use Committee (IACUC protocol # 10682). Target Site Blocker (TSB17) and scrambled control (SC) oligonucleotides were synthesized with full phosphorothioate backbones and 2′-O-methyl modifications (Integrated DNA Technologies). Mice received 5 mg/kg of TSB17 or SC oligonucleotide via intraperitoneal injection 16 hours prior to lipopolysaccharide (LPS) challenge, followed by a second dose at the time of LPS administration. Systemic inflammation was induced by intraperitoneal injection of LPS (Escherichia coli O111:B4, Sigma-Aldrich) at 5 mg/kg. Blood was collected by cardiac puncture 7 hours post-LPS injection, and serum was isolated by centrifugation and stored at −80°C until use. Serum concentrations of IL-17A and IL-6 were quantified using ELISA kits (R&D Systems, Minneapolis, MN) per manufacturer’s instructions and reported in pg/mL based on standard curves. Data were analyzed using GraphPad Prism 10.0 (GraphPad Software, San Diego, CA), and statistical significance was determined using one-way ANOVA followed by Tukey’s post-hoc test, with p-values < 0.05 considered significant.

### Experimental Autoimmune Uveitis (EAU) Induction and Treatment

#### Animals

Female Lewis rats (6–8 weeks old, 150–180 g; Charles River Laboratories) were housed under specific pathogen-free conditions with ad libitum access to food and water. All animal procedures were approved by the Institutional Animal Care and Use Committee (IACUC) and conducted in accordance with the ARVO Statement for the Use of Animals in Ophthalmic and Vision Research.

#### EAU Induction

EAU was induced by subcutaneous immunization at the base of the tail and flanks with a 200 μL emulsion containing 50 μg of bovine interphotoreceptor retinoid-binding protein peptide (IRBP₁₁₇₇₋₁₁₉₁) (Sigma-Aldrich) dissolved in PBS and emulsified 1:1 with Complete Freund’s Adjuvant (CFA) supplemented with 2.5 mg/mL Mycobacterium tuberculosis H37Ra (Difco). Rats were monitored daily for clinical signs of uveitis starting on Day 7 post-immunization.

#### Treatment Regimen

Rats were randomized into treatment groups (n = 5/group) and received intraperitoneal (IP) injections of TSB17 at 5 mg/kg once daily from Days 6 to 10 post-immunization. Additional groups received intravitreal (IVT) injections of 5 μL of oligonucleotides (40 μg) on Days 6 and 8. Dexamethasone (Sigma-Aldrich) was used as a positive control (40 μg/eye, IVT, Days 6 and 8). Vehicle-treated rats received either PBS or buffer alone.

#### Clinical Evaluatio

Fundoscopic examination was performed using a Micron IV retinal imaging system (Phoenix Research Labs) under isoflurane anesthesia on Days 0, 7, 9, 11, 13, and 14 post-immunization. Eyes were dilated with topical tropicamide and phenylephrine. Clinical EAU scores were assigned in a blinded fashion using a standard 0–4 grading scale:

- 0: No disease
- 1: Minimal vasculitis or optic disc swelling
- 2: Moderate vasculitis and infiltrates
- 3: Retinal hemorrhage or detachment
- 4: Complete retinal destruction or pan-uveitis

#### Histological Analysis

At the study endpoint (Day 14), animals were euthanized, and eyes were enucleated, fixed in 4% paraformaldehyde, paraffin-embedded, and sectioned (5 μm). Sections were stained with hematoxylin and eosin (H&E) for evaluation of retinal architecture and leukocyte infiltration. Histopathological scoring was performed as previously described (Ref), ranging from 0 (normal) to 4 (complete retinal destruction).

#### Statistical analysis

Data are shown as mean ± SD or SEM as indicated. Statistical analysis was performed using GraphPad Prism 10. Comparisons were made using: Student’s t-test, one-way ANOVA, two-way ANOVA with Tukey post-hoc test. AUC values were calculated where indicated. p < 0.05 was considered significant.

## ABBREVIATIONS

TSB: Target Site Blocker
RBP: RNA-binding protein
ARE: AU-rich element
cARE: conserved AU-rich element
RIP: RNA immunoprecipitation
SC: scrambled control oligonucleotide
miR-466: miR-466l-3p
3′UTR: untranslated region
MS2-TRAP: MS2-tagged RNA affinity purification
EAE: Experimental autoimmune encephalomyelitis
EAU: Experimental autoimmune uveitis
PMA: phorbol myristate acetate
PLL: poly-L-lysine
AUC: area under the curve

---

**TABLE 1.**
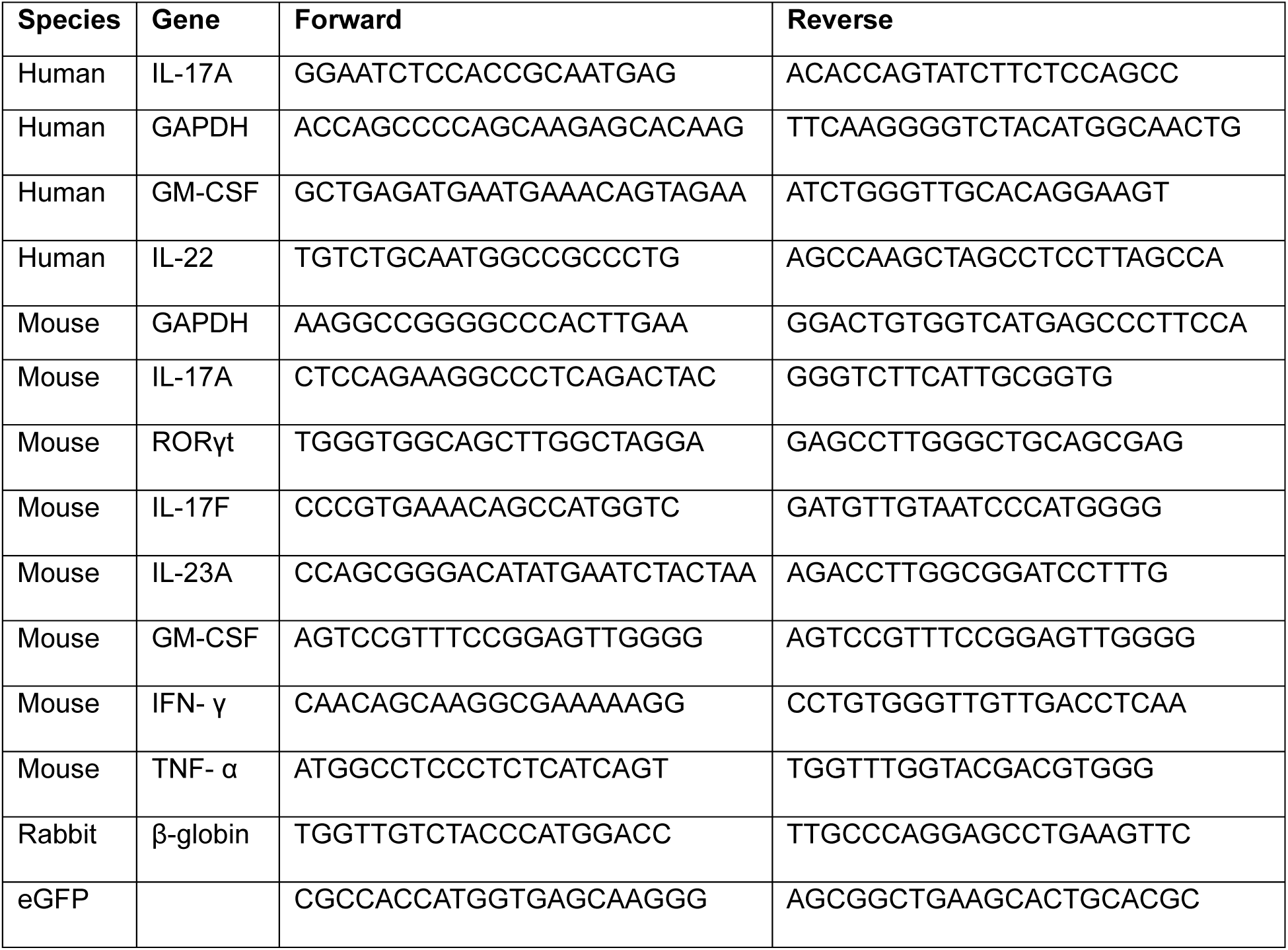
Primers for qRT-PCR.

**TABLE 2.**
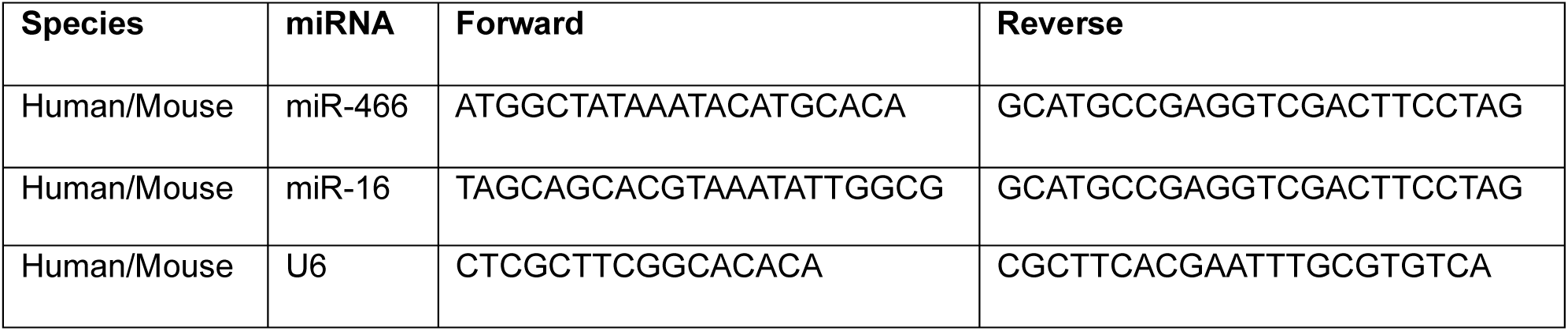
Primers for miRNA detection.

**TABLE 3.**
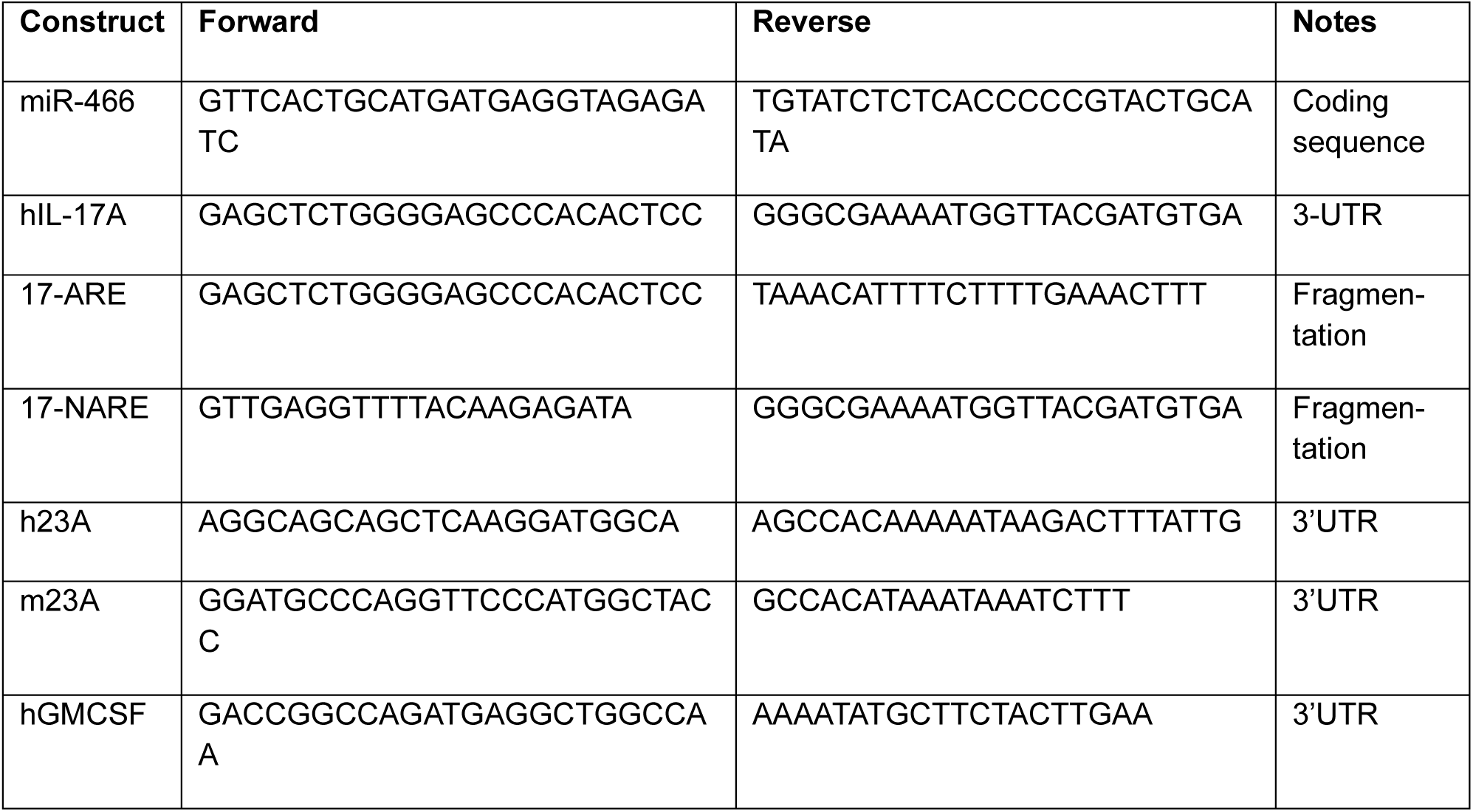
Primers used for cloning.

**TABLE 4.**
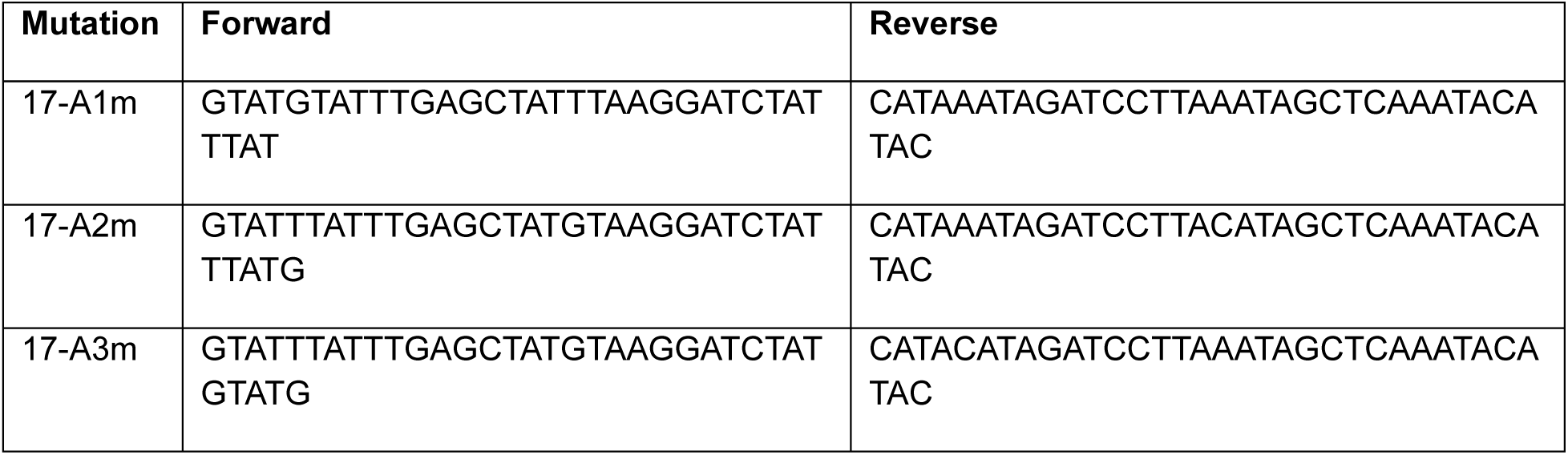
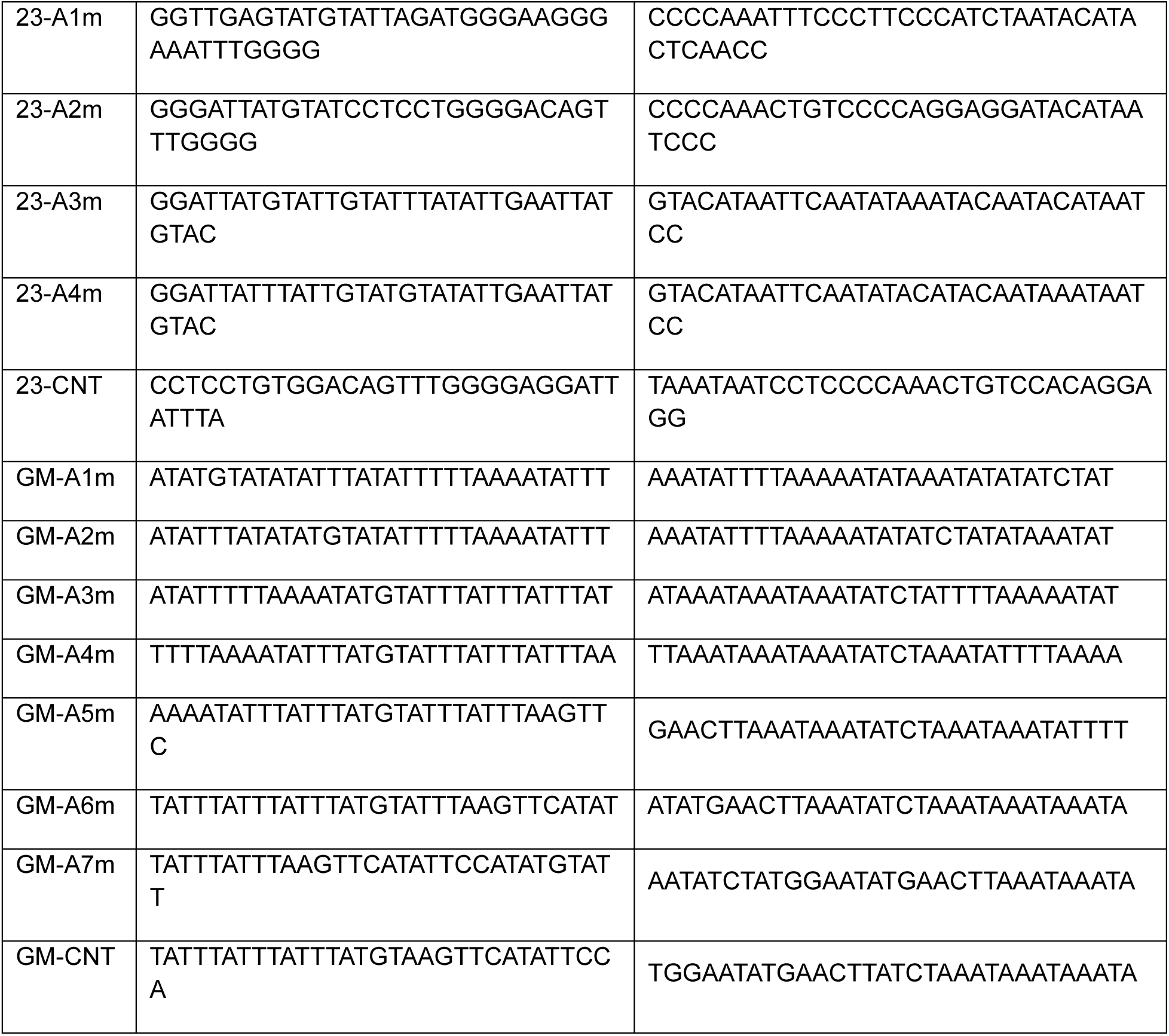
Primers for Site-directed mutagenesis.

**TABLE 5.**
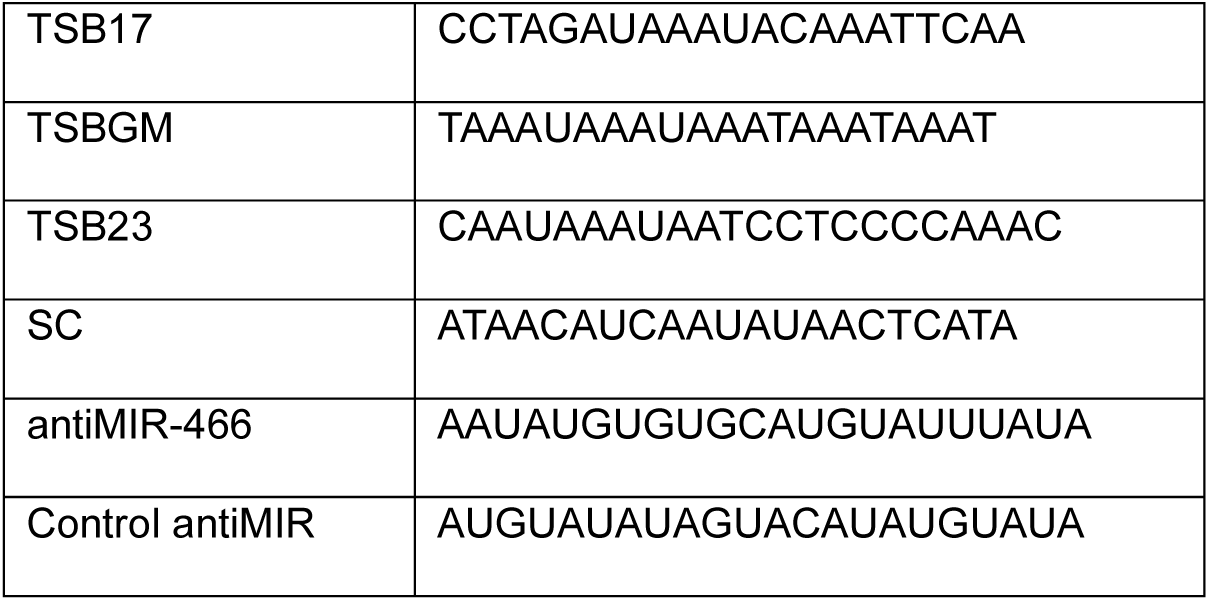
Sequences for oligonucleotides.

