## Supplementary material for "Disrupting miR-466l-3p and HuR Cooperation with Target Site Blockers Reveals a Therapeutic Strategy to Destabilize mRNA Transcripts": Suppl. Fig

### SUPPLEMENTAL FIGURES

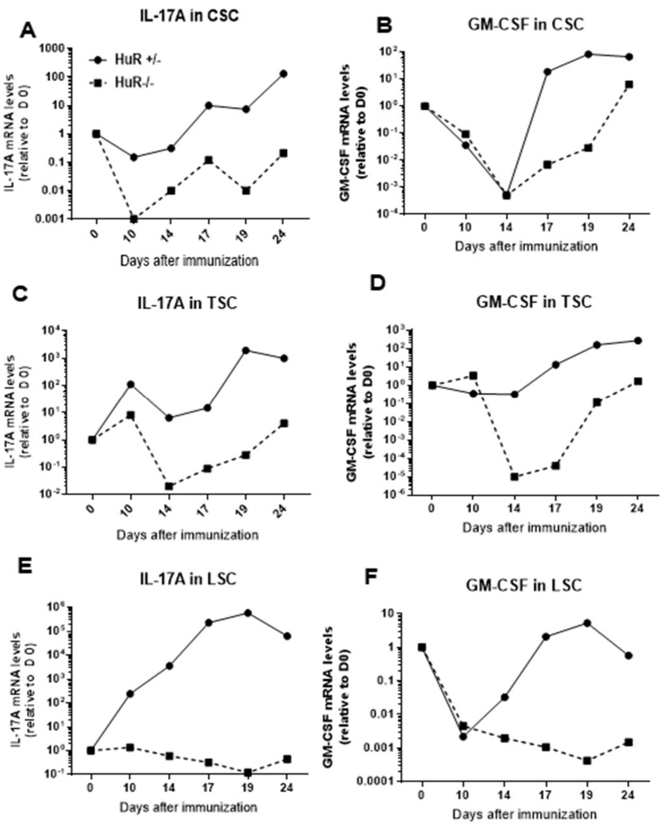


**Fig.S1: Post-Transcriptional Regulation of IL-17A and GM-CSF in the Spinal Cord of HuR-Deficient Mice During EAE**
EAE was induced in $HuR^{-}/^{-}$ and $HuR^{+}/^{-}$ mice via subcutaneous immunization with 200 µg $MOG_{35}$–$_{55}$ peptide emulsified in 250 µg complete Freund's adjuvant (CFA). Clinical symptoms were monitored daily. At days 0, 10, 14, 17, 19, and 24 post-immunization, one mouse per group was euthanized and spinal cords were isolated and sectioned into (CSC) (***A,B***), thoracic (TSC) (***C,D***), and lumbar (LSC) (**E,F**) regions. Total RNA was extracted, and mRNA levels of IL-17A and GM-CSF were quantified by qRT-PCR using gene-specific primers. Data are presented as fold change relative to naïve, unimmunized controls (day 0).

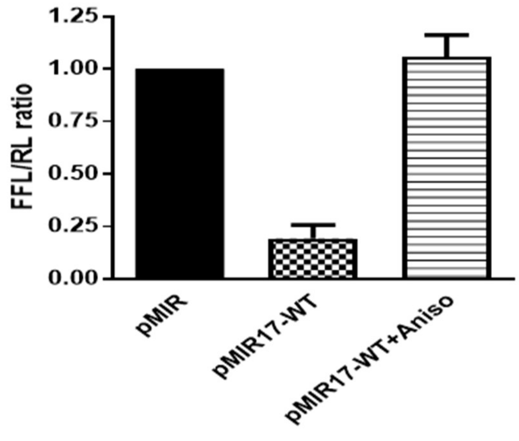


**Fig. S.2**: **Anisomycin treatment in dual-reporter assay**
NIH-3T3 cells were transfected with dual-luciferase pMIRGlo reporter plasmids containing either the human IL-17A 3′UTR (pMIR-h17) or the empty vector (pMIRGlo) for 24 hours. Cells were then either left untreated or stimulated with 1 µg/mL anisomycin for 6 hours. Following treatment, cells were lysed and Firefly (FF) and Renilla (RL) luciferase activities were measured. Firefly luciferase levels were normalized to Renilla luciferase, and fold induction was calculated relative to the pMIRGlo. Data is the average ±SEM of 3 independent experiments.

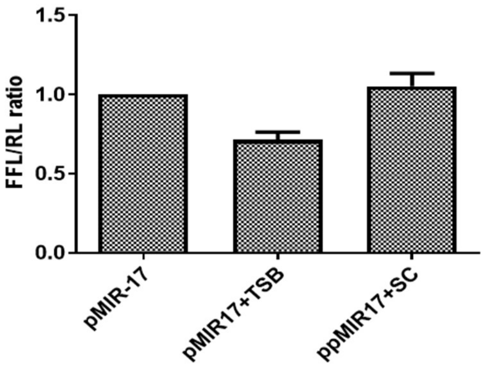


**Fig. S.3**: **TSB17 in a Dual-Luciferase Assay** Approximately 0.25.$10^6$ million NIH-3T3 cells were transfected with the dual-luciferase pMIRGlo reporter plasmid containing the wild-type IL-17A 3′UTR (pMIR-17A-WT) in the presence of 25 nM TSB17 or SC oligonucleotides. After 24 hours, firefly and Renilla luciferase activities were measured. Firefly luciferase levels were normalized to Renilla luciferase and expressed as fold change relative to empty vector controls. Data is the average ±SEM of 3 independent experiments.

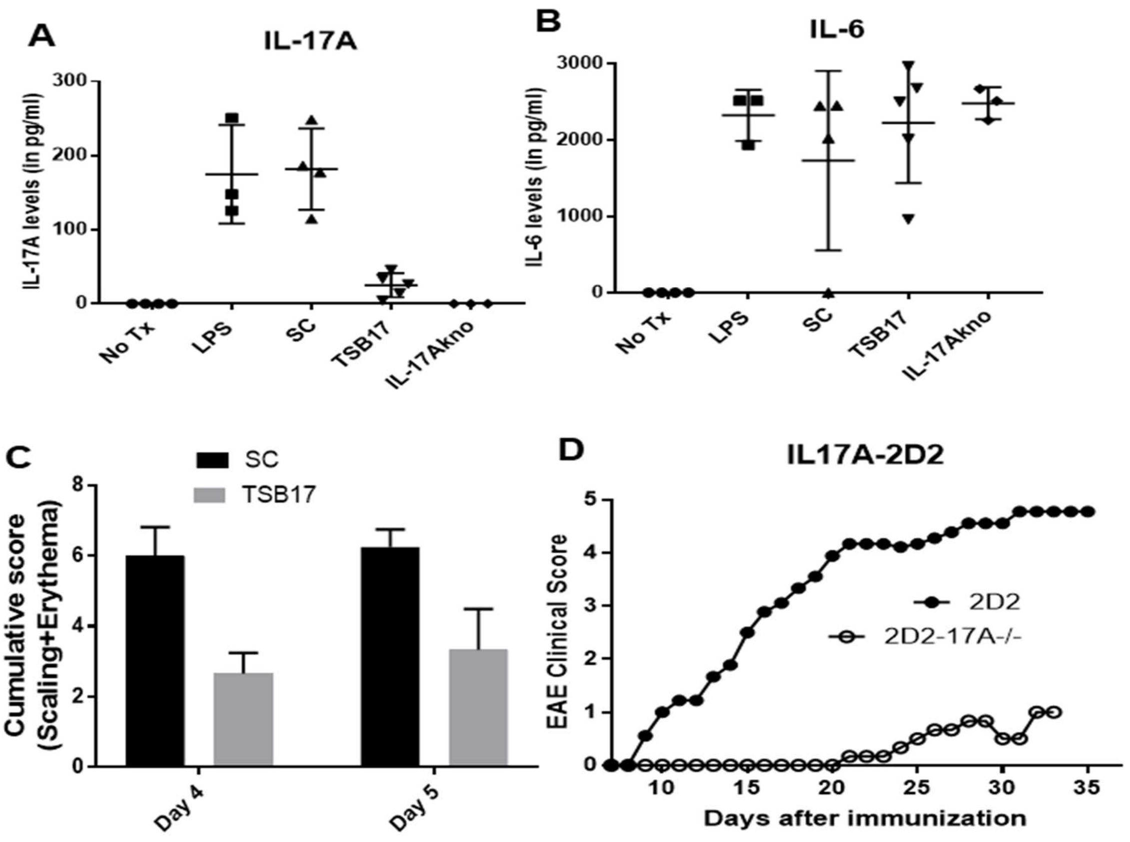


**Fig. S4: TSB17 attenuates IL-17A–mediated inflammation across multiple in vivo models** (***A/B***) Mice were administered 5 mg/kg TSB17 or SC oligonucleotide intraperitoneally 16 hours before and at the time of 5 mg/kg intraperitoneal injection with LPS as follow: LPS alone (n = 3), TSB17 (n = 5), SC (n = 4), IL-17A knockout mice (n=3), or naive controls (n=3). Mice were euthanized after 7 hours after LPS administration and sera was collected. (***A***) Serum IL-17A levels. mice. (***B***) Serum IL-6 levels. Data represent mean ± SEM. (***C***) C57BL/6 and IL-17A$^{-/-}$ mice were treated with daily imiquimod. C57BL/6 mice also received either SC or TSB17 oligonucleotide in 10% Pluronic F-127, applied topically 8 hours post-imiquimod. Scaling and erythema according to the PASI scoring criteria. (***D***) EAE was induced in 2D2-TCR (n=9) and 2D2-TCR-IL-17A$^{-/-}$ (n=6) transgenic mice using subcutaneous immunization at two flanking sites with 250 ug $MOG_{35-55}$ peptide and 500 ug zymosan adjuvant. Clinical symptoms were monitored and scored daily.

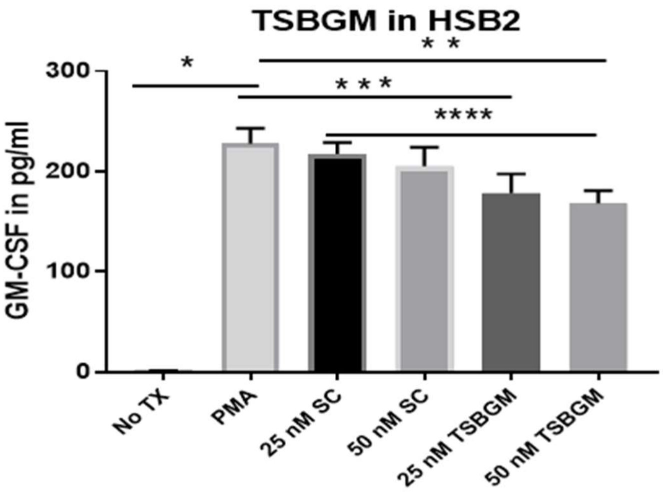


**Fig. S5: TSB-GM reduces GM-CSF protein expression and disrupts HuR binding to GM-CSF mRNA** (***A***) One million HSB-2 cells were transfected with either a SC oligonucleotide or increasing concentrations (25, 50, and 100 nM) with TSB-GM, followed by stimulation with 2ng/ml PMA for 24 hrs. GM-CSF levels in the culture supernatant were measured by ELISA 24 hours post-stimulation. Data represent mean ± SEM from three independent experiments. *$p < 0.05$ by one-way ANOVA with post hoc test.
